# Integrated Proteo-Metabolomics of Urinary Extracellular Vesicles Reveals Early Molecular Divergence and Temporal Pathogenesis of Sepsis-Associated AKI

**DOI:** 10.64898/2026.07.23.740423

**Authors:** Tanyu Chang, I-Lin Tsai, Guan-Yuan Chen, Te-I Weng, San-Yuan Wang, Yueh-Chu Sio, Ching-Yi Chen, Li-Yee Hong, I-Jen Chiu, Yen-Chung Lin, Hsi-Hsien Chen, Wei-Chiao Chang, Mai-Szu Wu, Michael X. Chen, Chih-Chin Kao

## Abstract

**Background:** Sepsis-associated acute kidney injury (S-AKI) is a major contributor to morbidity and mortality in critically ill patients. However, the molecular mechanisms underlying its temporal progression remain poorly understood because conventional biomarkers primarily reflect renal dysfunction rather than disease pathogenesis. Urinary extracellular vesicles (uEVs), which carry kidney-derived molecular cargo, provide a promising platform for monitoring renal-specific biological alterations during disease progression.

**Methods:** We conducted a longitudinal multi-omics study of uEVs collected from 81 patients with sepsis, including 48 patients with S-AKI and 33 sepsis-only controls. Patients were randomly assigned to a discovery cohort (n = 52) and an independent validation cohort (n = 29). Urine samples were collected at Day 1, Day 4, and Day 8 after AKI diagnosis. High-resolution proteomic and metabolomic profiling was performed to characterize temporal molecular alterations. Enriched pathways identified in the discovery cohort were evaluated in the validation cohort using pathway-level concordance analysis.

**Results:** Comparative analysis between S-AKI and sepsis-only patients identified distinct stage-specific molecular alterations throughout disease progression. At the early stage (Day 1), validated pathways included complement and coagulation cascades, ferroptosis, HIF-1 signaling, sphingolipid metabolism, and arachidonic acid metabolism, highlighting coordinated inflammatory, hypoxic, and lipid metabolic responses. During the mid-stage (Day 4), persistent activation of complement and coagulation cascades, ferroptosis, and HIF-1 signaling was accompanied by metabolic reprogramming involving alanine, aspartate and glutamate metabolism and tyrosine metabolism. Although limited sample availability reduced statistical power at Day 8, phenylalanine metabolism remained validated in the metabolomic analysis, suggesting persistent metabolic dysregulation during late-stage disease progression.

**Conclusions:** This study provides the first longitudinal, independently validated multi-omics characterization of human uEVs in S-AKI. By integrating proteomic and metabolomic profiling, we reveal the temporal evolution of renal-specific molecular pathways from early inflammatory and hypoxic responses to subsequent metabolic reprogramming. These findings establish uEV-based multi-omics as a promising strategy for molecular phenotyping of S-AKI beyond conventional clinical biomarkers and provide a valuable resource for future biomarker discovery and therapeutic target identification.

## Introduction

Sepsis-associated acute kidney injury (S-AKI) is a critical complication in intensive care units, accounting for nearly half of all AKI cases and carrying a 6- to 8-fold increase in hospital mortality [1]. Beyond acute survival, S-AKI triples the long-term risk of chronic kidney disease (CKD), imposing a significant global healthcare burden. Currently, clinical diagnosis relies on the KDIGO criteria—serum creatinine and urine output. However, these parameters are lagging functional indicators that reflect a decline in glomerular filtration rather than the specific molecular etiology of renal damage. Given the immense pathophysiological heterogeneity of S-AKI, there is an urgent need for molecular phenotyping tools that move beyond generic functional assessment toward precision medicine.

Recently, several functional and damage-specific biomarkers have been integrated into clinical research to identify AKI earlier than serum creatinine. Notable examples include Neutrophil Gelatinase-Associated Lipocalin (NGAL)[2, 3], Kidney Injury Molecule-1 (KIM-1)[4, 5], and the FDA-cleared cell-cycle arrest markers [TIMP-2]·[IGFBP7] (NephroCheck)[6, 7]. Additionally, newer candidates such as Proenkephalin A (penKid)[8, 9], a surrogate for real-time glomerular filtration rate (GFR) assessment, and L-FABP, a marker of tubular oxidative stress, have shown promise in various acute settings [10, 11]. However, the diagnostic and prognostic performance of these markers in the intensive care unit (ICU) is frequently compromised by the systemic complexity and high heterogeneity of sepsis. The primary limitation remains the susceptibility to systemic inflammatory “noise.” NGAL, for instance, is an acute-phase reactant heavily secreted by activated neutrophils during the systemic inflammatory storm; thus, its elevation in septic patients often reflects systemic inflammation rather than intrinsic renal-specific injury [12]. Similarly, while [TIMP-2]·[IGFBP7] effectively signals G1 cell-cycle arrest as a distress response, studies indicate that systemic inflammatory stress can trigger these arrest signals even in the absence of established structural kidney damage, potentially leading to over-diagnosis and poor specificity for S-AKI [13]. Furthermore, these traditional markers are largely unidimensional, acting as binary “stress signals” that fail to capture the complex, multi-pathway molecular reprogramming—such as ferroptosis, microvascular coagulation, or innate immune activation—that defines the pathogenesis of S-AKI [14]. While functional markers like penKid offer improved kinetic assessment of renal function, they provide limited insight into the underlying parenchymal pathophysiology. Therefore, there is a critical need for an organ-specific biological window, such as uEV multi-omics, that is shielded from systemic interference and provides a multidimensional landscape of the renal microenvironment to enable precise molecular phenotyping.

Extracellular vesicles (EVs), including exosomes and microvesicles, are nano-sized, membrane-bound particles released by cells to facilitate intercellular communication through the transport of bioactive proteins, lipids, and metabolites[15]. In the urinary tract, these vesicles are directly shed from renal tubular cells and podocytes, offering a non-invasive “liquid biopsy” of the renal microenvironment [16, 17]. Crucially, the lipid bilayer of uEVs encapsulates and protects their molecular cargo from enzymatic degradation[18], preserving proteins and nucleic acids that faithfully reflect the physiological and pathological status of their renal cells of origin, thereby providing kidney-specific molecular information that is less influenced by systemic circulation. While previous “bulk” urinary studies have provided foundational insights, they often struggle to isolate these protected, organ-specific signatures. Furthermore, existing uEV research has been largely limited by cross-sectional designs or single-omic perspectives, which fail to capture the longitudinal molecular evolution and the dynamic metabolic-proteomic interplay that drives S-AKI progression [19].

In this study, we conducted a comprehensive longitudinal multi-omics investigation of uEVs to decipher the temporal molecular landscape of S-AKI within the complex ICU environment. To distinguish renal-specific molecular alterations from systemic inflammatory responses, we employed a rigorous comparative design using sepsis-only patients as controls and further evaluated our findings in an independent validation cohort. By integrating high-resolution proteomic and metabolomic profiling, we identified stage-specific molecular programs underlying S-AKI progression, revealing early activation of complement and coagulation cascades, ferroptosis, HIF-1 signaling, sphingolipid metabolism, and arachidonic acid metabolism, followed by metabolic reprogramming characterized by alterations in amino acid metabolism during disease progression. To our knowledge, this is the first longitudinal multi-omics study of human uEVs to systematically delineate the temporal molecular evolution of S-AKI and independently validate pathway-level alterations, providing a robust framework for molecular phenotyping beyond conventional clinical biomarkers.

## Methods

### Study design and patient enrollment

This was a longitudinal prospective study evaluating urinary EV proteomic and metabolomic profiles in patients with sepsis, with or without AKI. The overall study design is illustrated **(Figure 1)**. The sample size was determined based on the availability of well-characterized clinical samples. While no formal power calculation was conducted, the sample size was sufficient to identify significant expression differences with biological relevance. Patients with sepsis were recruited from the ICUs at Taipei Medical University Hospital (Taipei, Taiwan) between November 25, 2022, and January 3, 2024. The study was approved by the Taipei Medical University Joint Institutional Review Board (IRB No. N202404097), and informed consent was obtained from all participants. AKI was diagnosed based on the KDIGO criteria [20]. Eighty-one patients with sepsis were enrolled, including 48 with sepsis-associated acute kidney injury (S-AKI) and 33 sepsis-only controls. The patients were randomly assigned to a discovery cohort (n = 52) and a validation cohort (n = 29). Baseline demographic and clinical characteristics were collected for all participants. Proteomic and metabolomic analyses were performed on urinary extracellular vesicle (uEV) samples collected at three time points: Day 1 (D1; the day of AKI diagnosis), Day 4 (D4; 3 days after diagnosis), and Day 8 (D8; 7 days after diagnosis). The discovery cohort included 52, 44, and 22 samples at D1, D4, and D8, respectively, whereas the validation cohort included 29, 21, and 9 samples at the corresponding time points. For patients without AKI, the diagnostic time point (D-day) was defined as the date of sepsis diagnosis. Differentially expressed proteins (DEPs) and differentially expressed features (DEFs) consistently demonstrating significant differences at the three time points were identified as candidate biomarkers through Venn diagram analysis.

**Figure 1.**
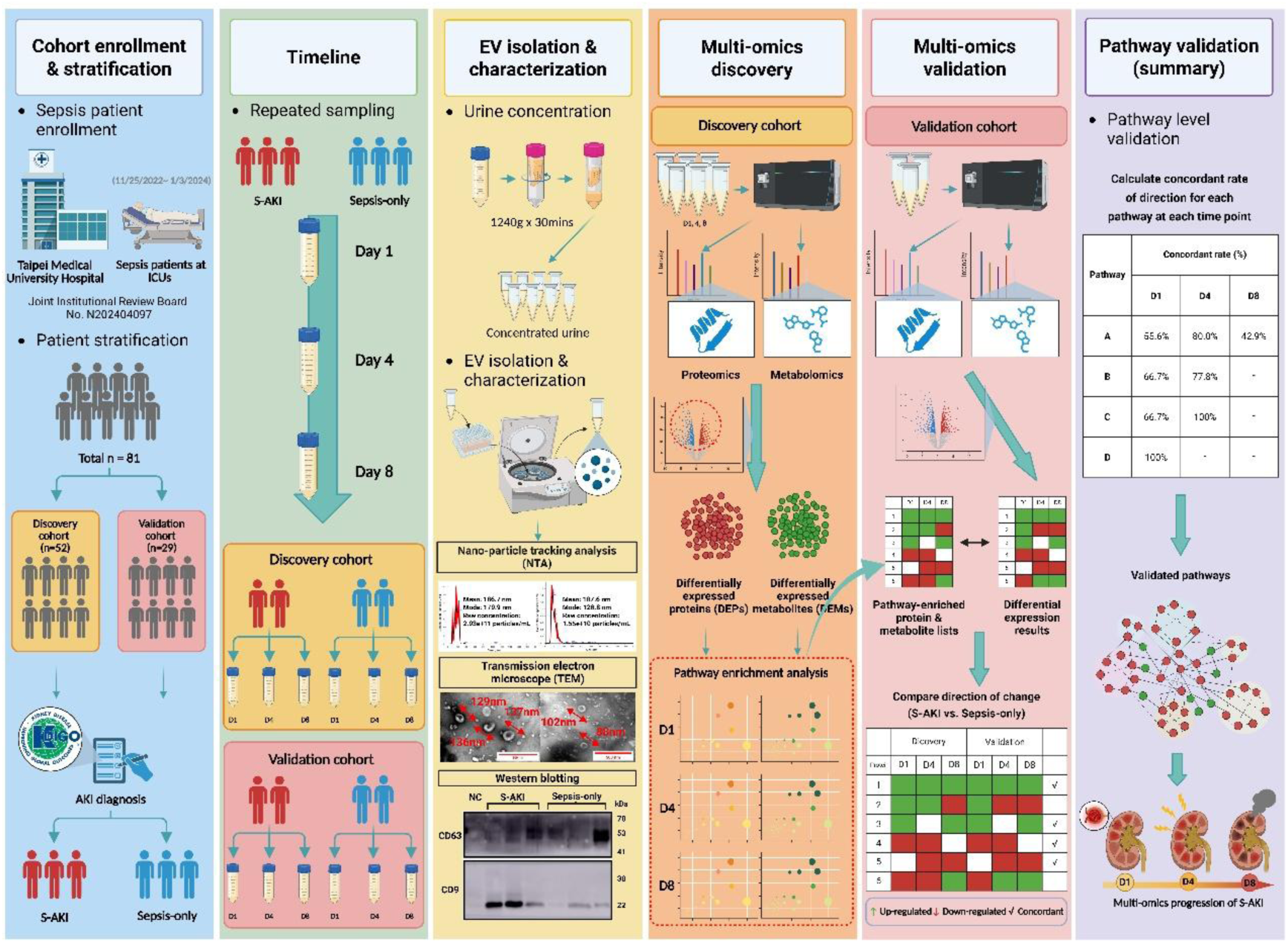
Study design and characterization of urine EVs. This figure illustrates the overall study workflow, including (1) cohort enrollment, (2) study timeline, (3) EV isolation and characterization, (4, 5) multi-omics profiling in the discovery cohort followed by validation in an independent cohort, and (6) pathway-level validation based on concordance analysis. EV characterization includes Nano-particle tracking analysis (NTA), transmission electron microscopy (TEM), and Western blotting results of urine EVs. CD63 (30-50 kDa), CD9 (24-30 kDa)

### EV isolation and characterization

Urinary EVs were isolated as outlined in **Figure 1**. Briefly, 15 mL of urine was concentrated to 500 µL using Amicon Ultra-15 centrifugal filters (100 kDa cutoff; Merck Millipore, Darmstadt, Germany). EVs were purified using the SmartSEC™ HT EV Isolation System (System Biosciences, Palo Alto, CA, USA), following the manufacturer’s instructions [21]. To confirm EV isolation, the following validation techniques were employed: first, western blotting for exosomal markers CD63 and CD9 (System Biosciences, Palo Alto, CA, USA) [22], with 20 μg of protein per sample was performed. Second, nanoparticle tracking analysis (NTA) using the NanoSight NS300 (Malvern Panalytical, UK) to assess particle size and concentration was done. Finally, transmission electron microscopy (TEM) using a Hitachi HT-7700 (Hitachi, Japan) to examine EV morphology was performed.

### Sample preparation for proteomic and metabolomic analysis

Proteins were precipitated with ice-cold acetone (Honeywell, Charlotte, NC, USA) at a volume four times that of each sample, and 50 µg of protein was processed per sample. Pellets were resuspended in 100 µL of 6 M urea (Nihon Shiyaku Industries, LTD, Japan), reduced with 2 µL of 550 mM dithiothreitol (DTT; Sigma-Aldrich, St. Louis, MO, USA), alkylated with 4 µL of 450 mM iodoacetamide (IAA; Sigma-Aldrich), and digested with 5 µL of trypsin (0.2 µg/µL; Promega, Madison, WI, USA). Solid-phase extraction (SPE) was performed using Sep-Pak® Vac 1cc (50mg) C18 cartridges (Waters, Milford, MA, USA) and a SpeedVac concentrator to purify and concentrate peptides. Meanwhile, the supernatant was collected for metabolomic analysis. Like proteomic analysis, metabolites were concentrated using a SpeedVac concentrator. The dried extracts were reconstituted in 50% methanol for LC–MS/MS analysis.

### Liquid chromatography-mass spectrometry (LC-MS)

For proteomic analysis, MS and gradient settings optimized in our previous work [23] were applied. Analyses were performed on an Orbitrap Fusion Lumos Tribrid quadrupole-ion trap-Orbitrap mass spectrometer (Thermo Fisher Scientific, CA, USA) coupled with an Ultimate 3000 NanoLC system (Thermo Fisher Scientific, Bremen, Germany). Peptides were separated on a C18 Acclaim PepMap NanoLC column (75 μm ID × 25 cm). A segmented gradient (2%–40% acetonitrile in 0.1% formic acid) was applied over 40 minutes at a flow rate of 300 nL/min.

A full MS scan was conducted, followed by high-energy collision-activated dissociation (HCD)-MS/MS analysis of the most intense ions within 3 s. Data were externally calibrated to maintain a mass accuracy of <5 ppm. MS1 scans were acquired at a resolution of 120,000 (m/z 200), followed by HCD-MS/MS scans (resolution: 15,000) with dynamic exclusion (60 s), a 1.4 Da isolation window, and a normalized collision energy of 32%. The raw data were deposited in the ProteomeXchange Consortium via PRIDE (dataset identifier: PXD063789).

For metabolomic analysis, data acquisition was performed on a SYNAPT XS mass spectrometer (Waters, MA, USA) equipped with an ACQUITY UPLC system (Waters, Milford, MA, USA). Metabolites were separated on a Poroshell 120 EC-C18 column (1.9 μm, 2.1 × 100 mm). MS1 scanning ranges from 50 Da to 1200 Da, with a capillary voltage maintained at 2.0 kV, a source temperature of 150 °C, a desolvation temperature of 400 °C, a cone voltage of 40 V, and a cone gas flow of 50 L/h. The cone gas flow is 50 L/hr. The survey scan starts at a mass of 50 Da and ends at 1000 Da, with an intensity threshold of 10,000 and the detection of up to 10 components.

### Data Processing and Bioinformatics Analysis

Protein identification and label-free quantification were performed using MaxQuant (v2.4.7.0) with a reviewed human FASTA database (UniProtKB Taxonomy ID: 9606; downloaded September 12, 2023). Data were filtered and normalized in Perseus (v2.0.11). Proteins with missing values in more than 60% of the samples were excluded from the analysis. Downstream analyses—hierarchical clustering, sPLS-DA, and volcano plots—were conducted in MetaboAnalyst 6.0. DEPs are defined as proteins with a Wilcoxon test p-value < 0.1 and fold change > 1.3.

Metabolite identification and label-free quantification were conducted using Progenesis QI with the Human Metabolome Database (Metabolite Structures, Version 5.0, downloaded August 20, 2025). The adducts were selected according to a previous report[24]. The searched adducts included [M+H]^+^, [M+2H]^2+^, [M+Na]^+^, [M+K]^+^, and [M+NH₄]^+^ in positive-ion mode, and [M−H]^−^, [M−2H]^2−^, [M−2H+Na]^−^, and [M−2H+K]^−^ in negative-ion mode. Features with missing values in more than 60% of the samples were excluded from further analysis. Downstream analyses, hierarchical clustering, sPLS-DA, and volcano plots were also conducted in MetaboAnalyst 6.0. DEFs were defined as features with a Wilcoxon test p-value < 0.05 and fold change > 1.3. Metabolite annotation was conducted using a non-targeted workflow and achieved MSI level-3 confidence according to the Metabolomics Standards Initiative (MSI) guidelines[25]. Further validation using authentic reference standards and targeted MS/MS analyses will be required to confirm metabolite identities.

Finally, pathway analyses of the DEPs and DEMs were performed using MetaboAnalyst 6.0. Enriched pathway was identified based on p value (<0.05) and pathway impact (>0.05) to ensure the robustness of our interpretation. The enriched proteins and metabolites identified in the discovery cohort were evaluated in the independent validation cohort by assessing the concordance of fold-change directions between the two cohorts. A protein or metabolite was considered concordant when the direction of fold change (up- or down-regulation in S-AKI relative to the Sepsis-only group) was consistent between the discovery and validation cohorts at the corresponding time point. The pathway concordance rate was then calculated as the proportion of concordant proteins or metabolites among all evaluable proteins or metabolites within each enriched pathway. Pathways were considered validated when the concordance rate was ≥ 50% and at least two evaluable proteins or metabolites were available for assessment.

## Results

### Patients’ demographics and outcomes

A total of 52 patients with sepsis were enrolled in the study, including 29 patients in the S-AKI group and 23 in the sepsis-only control group. Patient characteristics are summarized in **Table 1**. The mean age was 69 years, and 28 (53.8%) were male. Baseline characteristics were generally comparable between the groups, except for differences in renal function and Sequential Organ Failure Assessment (SOFA) scores, which were significantly higher in the S-AKI group. The Acute Physiology and Chronic Health Evaluation II (APACHE II) score did not differ significantly between groups.

**Table 1.**
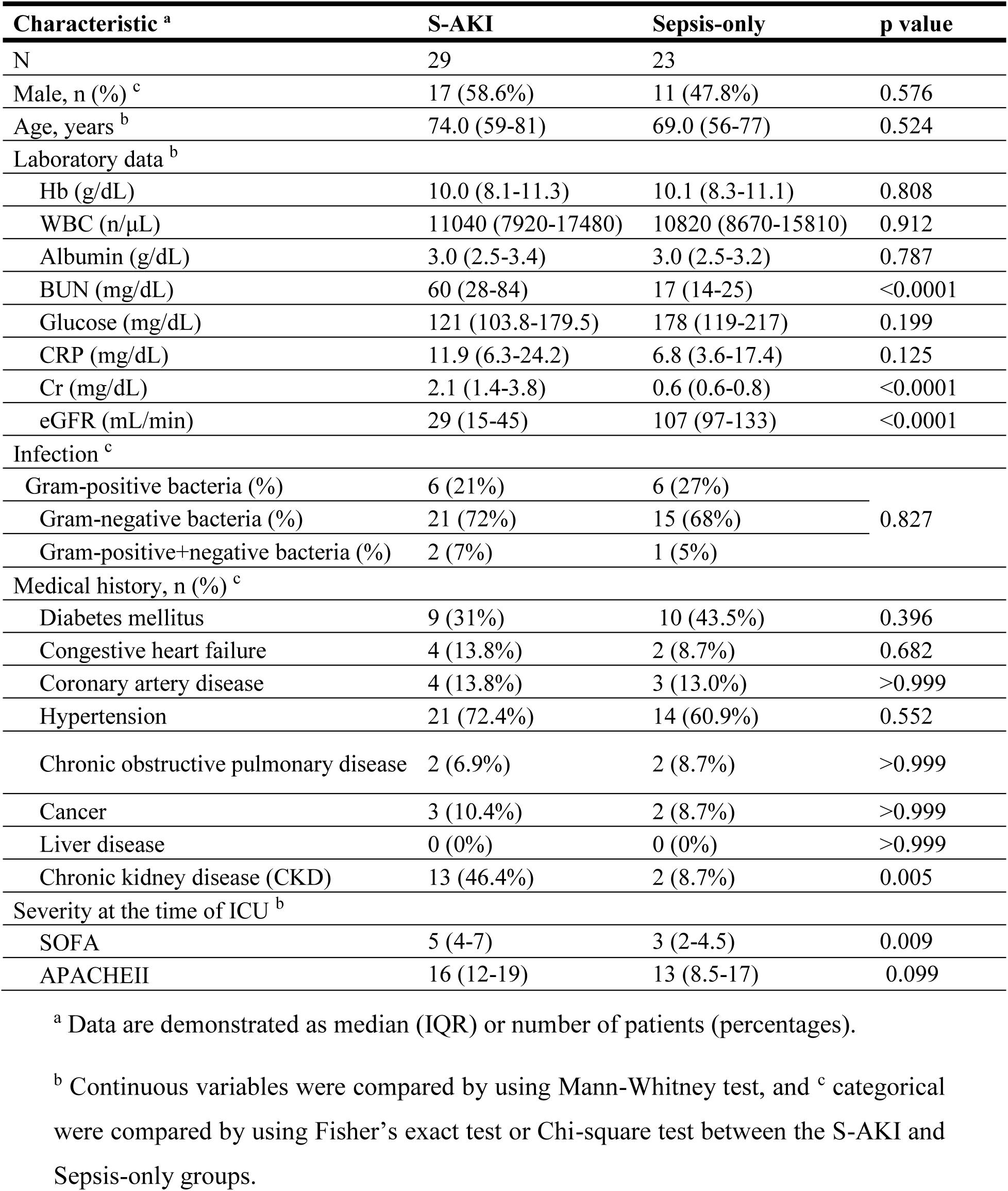
Baseline characteristics of S-AKI and sepsis-only patients.

| Characteristic <sup>a</sup> | S-AKI | Sepsis-only | p value |
| --- | --- | --- | --- |
| N | 29 | 23 |  |
| Male, n (%) <sup>c</sup> | 17 (58.6%) | 11 (47.8%) | 0.576 |
| Age, years <sup>b</sup> | 74.0 (59-81) | 69.0 (56-77) | 0.524 |
| Laboratory data <sup>b</sup> |  |  |  |
| Hb (g/dL) | 10.0 (8.1-11.3) | 10.1 (8.3-11.1) | 0.808 |
| WBC (n/μL) | 11040 (7920-17480) | 10820 (8670-15810) | 0.912 |
| Albumin (g/dL) | 3.0 (2.5-3.4) | 3.0 (2.5-3.2) | 0.787 |
| BUN (mg/dL) | 60 (28-84) | 17 (14-25) | <0.0001 |
| Glucose (mg/dL) | 121 (103.8-179.5) | 178 (119-217) | 0.199 |
| CRP (mg/dL) | 11.9 (6.3-24.2) | 6.8 (3.6-17.4) | 0.125 |
| Cr (mg/dL) | 2.1 (1.4-3.8) | 0.6 (0.6-0.8) | <0.0001 |
| eGFR (mL/min) | 29 (15-45) | 107 (97-133) | <0.0001 |
| Infection <sup>c</sup> |  |  |  |
| Gram-positive bacteria (%) | 6 (21%) | 6 (27%) | 0.827 |
| Gram-negative bacteria (%) | 21 (72%) | 15 (68%) |  |
| Gram-positive+negative bacteria (%) | 2 (7%) | 1 (5%) |  |
| Medical history, n (%) <sup>c</sup> |  |  |  |
| Diabetes mellitus | 9 (31%) | 10 (43.5%) | 0.396 |
| Congestive heart failure | 4 (13.8%) | 2 (8.7%) | 0.682 |
| Coronary artery disease | 4 (13.8%) | 3 (13.0%) | >0.999 |
| Hypertension | 21 (72.4%) | 14 (60.9%) | 0.552 |
| Chronic obstructive pulmonary disease | 2 (6.9%) | 2 (8.7%) | >0.999 |
| Cancer | 3 (10.4%) | 2 (8.7%) | >0.999 |
| Liver disease | 0 (0%) | 0 (0%) | >0.999 |
| Chronic kidney disease (CKD) | 13 (46.4%) | 2 (8.7%) | 0.005 |
| Severity at the time of ICU <sup>b</sup> |  |  |  |
| SOFA | 5 (4-7) | 3 (2-4.5) | 0.009 |
| APACHEII | 16 (12-19) | 13 (8.5-17) | 0.099 |

<sup>a</sup> Data are demonstrated as median (IQR) or number of patients (percentages).
<sup>b</sup> Continuous variables were compared by using Mann-Whitney test, and <sup>c</sup> categorical were compared by using Fisher's exact test or Chi-square test between the S-AKI and Sepsis-only groups.

During hospitalization, 5 patients (9.6%) required dialysis, and 10 (19.2%) died. At 90 days, the mortality rate was 21.1% (n = 11). Both in-hospital and 90-day mortality rates were significantly higher in the S-AKI group compared to the sepsis-only group **(Table 2)**.

**Table 2.** Outcome of S-AKI and sepsis-only patients.

| **Characteristic <sup>a</sup>** | **S-AKI** | **Sepsis-only** | **p value** |
| --- | --- | --- | --- |
| Need for dialysis, n (%) | 5 (17,2%) | 0 (0%) | 0.058 |
| In-hospital mortality, n (%) | 9 (31%) | 1 (4%) | 0.003 |
| 90-day mortality, n (%) | 10 (34%) | 1 (4%) | 0.014 |

<sup>a</sup> Data are presented as the number of patients (percentage). Categorical variables were compared between the S-AKI and sepsis-only groups using Fisher's exact test.

### Characterization of urinary extracellular vesicles

Isolated urinary EVs displayed a heterogeneous size distribution and exhibited the characteristic cup-shaped morphology, consistent with previous reports [15]. Western blot analysis confirmed the presence of exosomal markers CD63 and CD9 in both groups **(Figure 1)**.

### Identification of divergence in S-AKI and Sepsis-only groups on Day 1

sPLS-DA score plots demonstrated separation between the S-AKI and sepsis-only groups on Day 1 in both proteomic and metabolomic datasets, with component 1 explaining 6.5%, 4.9%, and 2.5% of the variance, respectively **(Figure 2A–C)**. Volcano plot analysis identified 50 differentially expressed proteins (DEPs), including 32 upregulated and 18 downregulated proteins in the S-AKI group, as well as 518 differentially expressed features (DEFs) in positive ion mode (202 upregulated and 316 downregulated) and 162 DEFs in negative ion mode (50 upregulated and 112 downregulated) **(Figure 2D-F)**.

**Figure 2.**
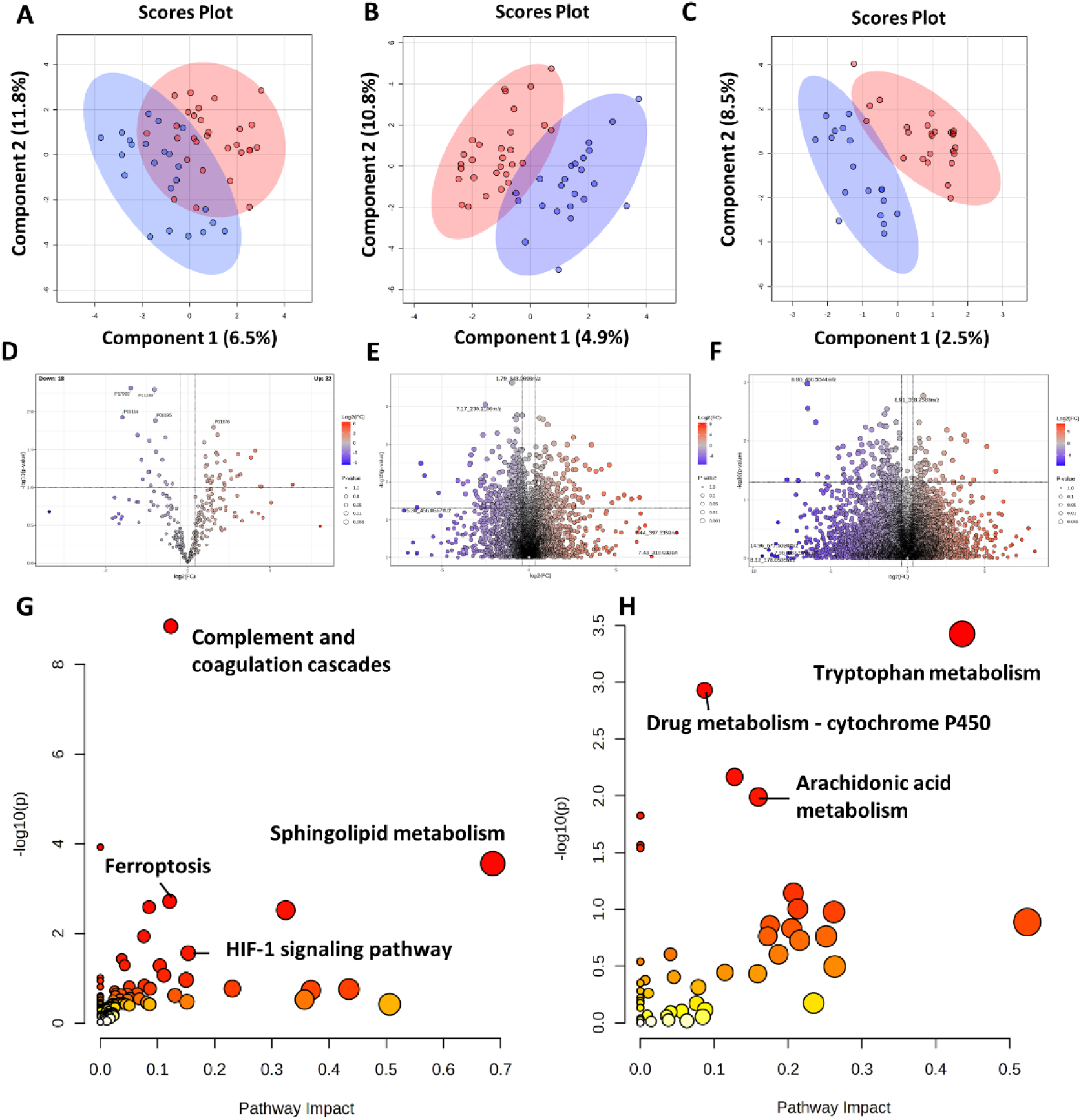
Proteomic and metabolomic analyses of the S-AKI and Sepsis-only groups on Day 1. A.–C. sPLS-DA score plots showing the separation between the S-AKI and sepsis-only groups based on proteomics and metabolomics datasets acquired in positive- and negative-ion modes on Day 1. D.–F. Volcano plots of proteomics and metabolomics datasets (positive- and negative-ion modes) on Day 1, illustrating statistical significance and fold change (FC). Red and blue dots indicate upregulated and downregulated molecules, respectively. Differentially expressed proteins/metabolites (DEPs/DEMs) were identified using a p value < 0.05 and FC > 1.3. G., H. Pathway enrichment analysis of differentially expressed proteins and metabolites from the merged metabolomics datasets on Day1 (positive- and negative-ion modes). DEPs, differentially expressed proteins; DEMs, differentially expressed metabolites; sPLS-DA, sparse partial least squares discriminant analysis; S-AKI, sepsis-induced acute kidney injury.

Pathway enrichment analysis revealed distinct biological alterations in both proteomic and metabolomic profiles **(Figure 2G, H)**. Collectively, these findings demonstrated significant molecular divergence between the S-AKI and sepsis-only groups at both the proteomic and metabolomic levels. Accordingly, the same analytical workflow, including volcano plot and pathway enrichment analyses, was subsequently applied to the Day 4 and Day 8 datasets **(Figures 3 and 4)**.

**Figure 3.**
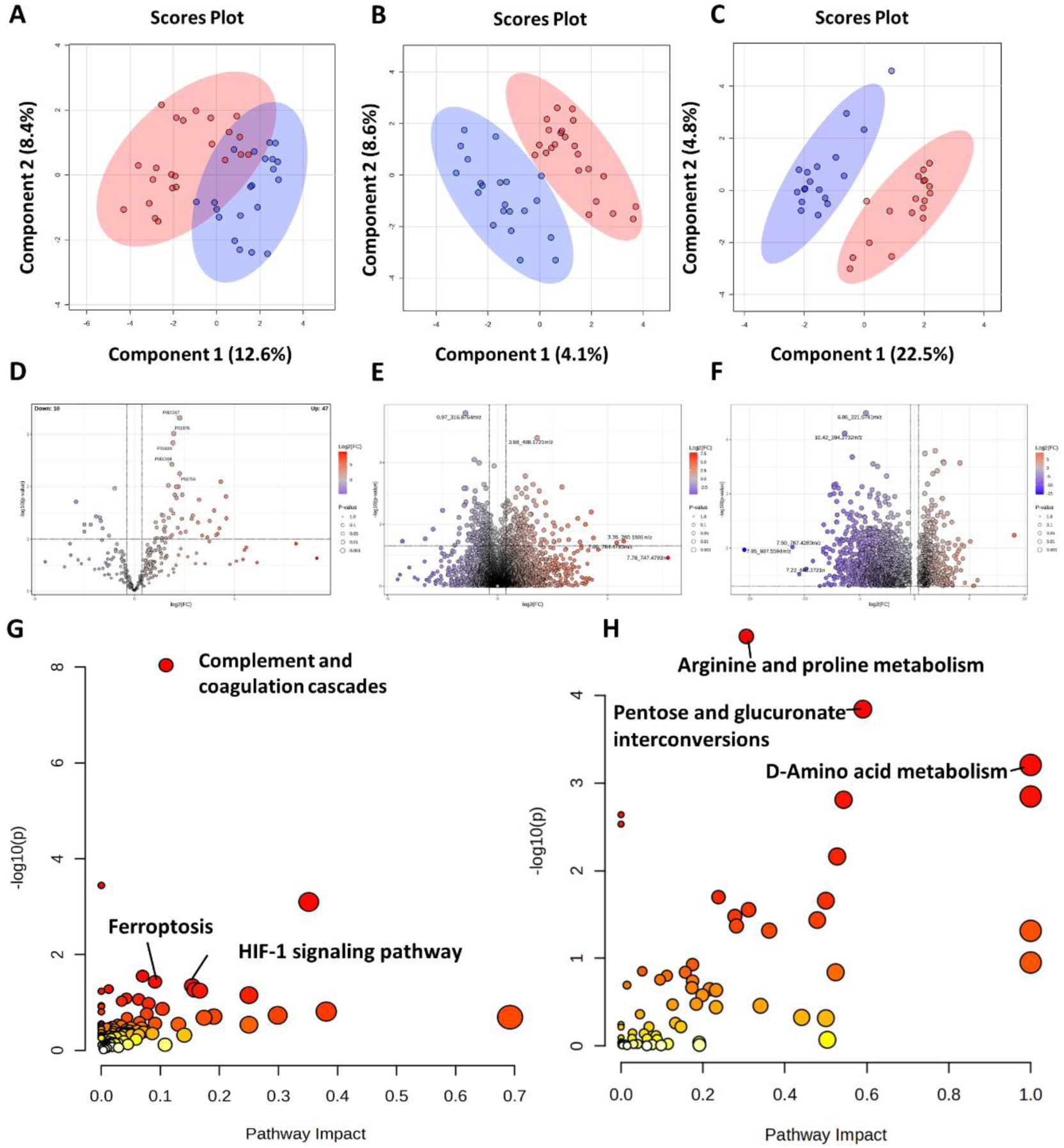
Proteomic and metabolomic analyses of the S-AKI and Sepsis-only groups on Day 4. A.–C. sPLS-DA score plots showing the separation between the S-AKI and sepsis-only groups based on proteomics and metabolomics datasets acquired in positive- and negative-ion modes on Day 4. D.–F. Volcano plots of proteomics and metabolomics datasets (positive- and negative-ion modes) on Day 4, illustrating statistical significance and fold change (FC). Red and blue dots indicate upregulated and downregulated molecules, respectively. Differentially expressed proteins/metabolites (DEPs/DEMs) were identified using a p value < 0.05 and FC > 1.3. G., H. Pathway enrichment analysis of differentially expressed proteins and metabolites from the merged metabolomics datasets on Day 4 (positive- and negative-ion modes). DEPs, differentially expressed proteins; DEMs, differentially expressed metabolites; sPLS-DA, sparse partial least squares discriminant analysis; S-AKI, sepsis-induced acute kidney injury.

**Figure 4.**
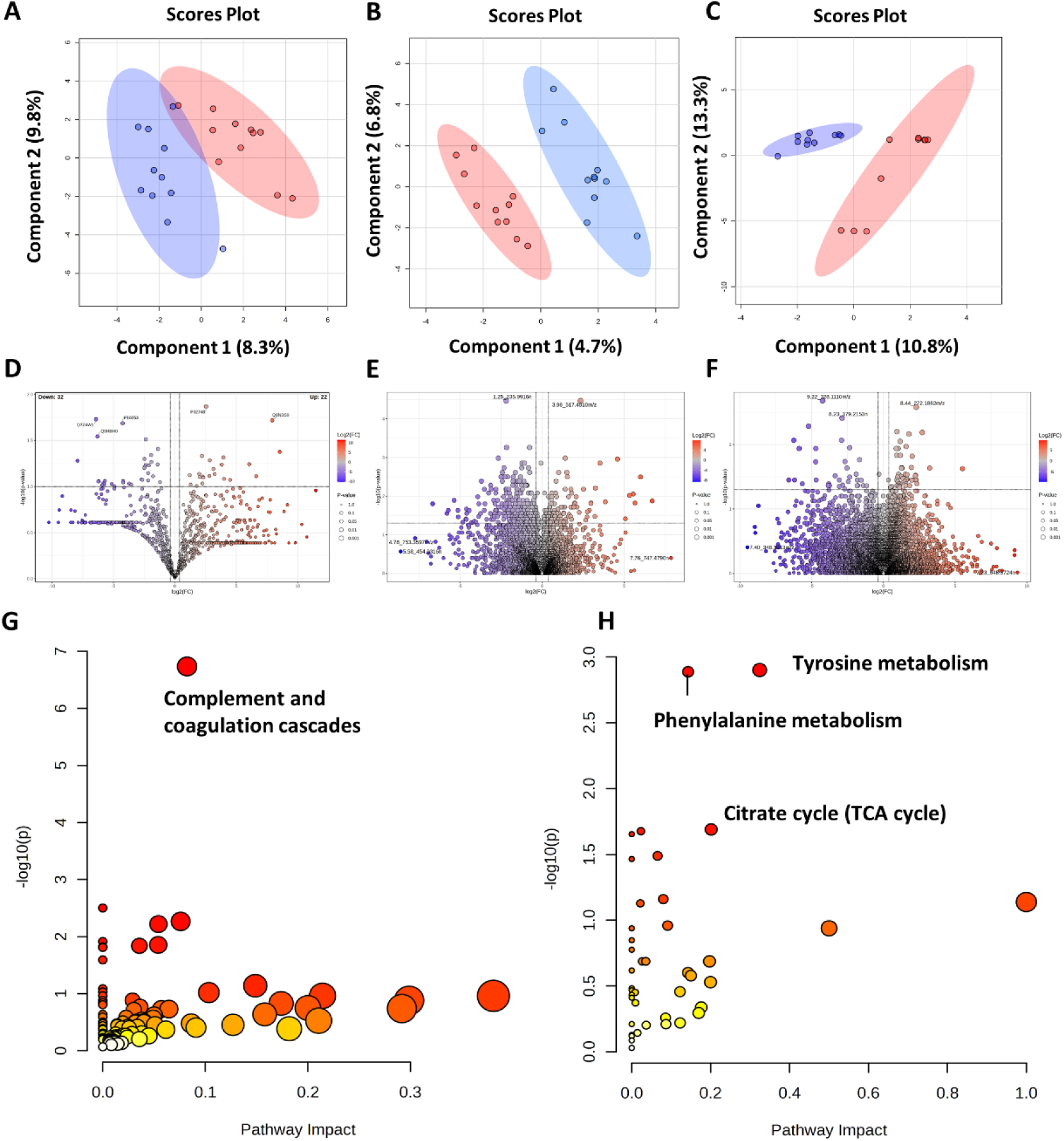
Proteomic and metabolomic analyses of the S-AKI and Sepsis-only groups on Day 8. A.–C. sPLS-DA score plots showing the separation between the S-AKI and sepsis-only groups based on proteomics and metabolomics datasets acquired in positive- and negative-ion modes on Day 8. D.–F. Volcano plots of proteomics and metabolomics datasets (positive- and negative-ion modes) on Day 8, illustrating statistical significance and fold change (FC). Red and blue dots indicate upregulated and downregulated molecules, respectively. Differentially expressed proteins/metabolites (DEPs/DEMs) were identified using a p value < 0.05 and FC > 1.3. G., H. Pathway enrichment analysis of differentially expressed proteins and metabolites from the merged metabolomics datasets on Day 8 (positive- and negative-ion modes). DEPs, differentially expressed proteins; DEMs, differentially expressed metabolites; sPLS-DA, sparse partial least squares discriminant analysis; S-AKI, sepsis-induced acute kidney injury.

### Proteomic and metabolomic validation at the pathway level

In the proteomic dataset, the validated pathways at D1 were sphingolipid metabolism, HIF-1 signaling pathway, complement and coagulation cascades, and ferroptosis, with concordance rates of 50.0%, 66.7%, 66.7%, and 66.7%, respectively. At D4, the HIF-1 signaling pathway, ferroptosis, and complement and coagulation cascades remained validated, showing concordance rates of 100.0%, 100.0%, and 77.8%, respectively. By D8, complement and coagulation cascades remained consistently enriched throughout disease progression, with a concordance rate of 42.9%, indicating partial agreement between the discovery and validation cohorts. **(Table 3)**. The validation results for all pathway-enriched proteins are summarized in **Table S1**.

**Table 3.** Summary of pathway validation between the Discovery and Validation cohorts of the proteomic dataset.

| Pathway | D1<br>Concordant<br>/ Total | D1<br>Validation<br>Rate (%) | D4<br>Concordant<br>/ Total | D4<br>Validation<br>Rate (%) | D8<br>Concordant<br>/ Total | D8<br>Validation<br>Rate (%) |
| --- | --- | --- | --- | --- | --- | --- |
| Complement<br>and<br>coagulation<br>cascades | 6/9 | 66.7 | 7/9 | 77.8 | 3/7 | 42.9 |
| Sphingolipid<br>metabolism | 2/4 | 50.0 | – | – | – | – |
| HIF-1<br>signaling<br>pathway | 2/3 | 66.7 | 3/3 | 100.0 | – | – |
| Ferroptosis | 2/3 | 66.7 | 2/2 | 100.0 | – | – |

Validation rates were calculated separately for each time point based on the proportion of proteins exhibiting concordant fold-change directions between the Discovery and Validation cohorts. Overall validation rates were calculated using all eligible protein–time point comparisons within each pathway. Concordance was defined as consistent up- or down-regulation in S-AKI relative to Sepsis-only between cohorts.

In the metabolomic dataset, the validated pathways at D1 were arachidonic acid metabolism and tryptophan metabolism, with concordance rates of 100.0% and 50.0%, respectively. At D4, caffeine metabolism, alanine, aspartate and glutamate metabolism, and tyrosine metabolism were validated, with concordance rates of 100.0%, 83.3%, and 50.0%, respectively. At D8, despite the limited sample size in the validation cohort, phenylalanine metabolism met the predefined validation criteria, with a concordance rate of 66.7%. **(Table 4)**. The validation results for all pathway-enriched metabolites are summarized in **Table S2**.

**Table 4.**
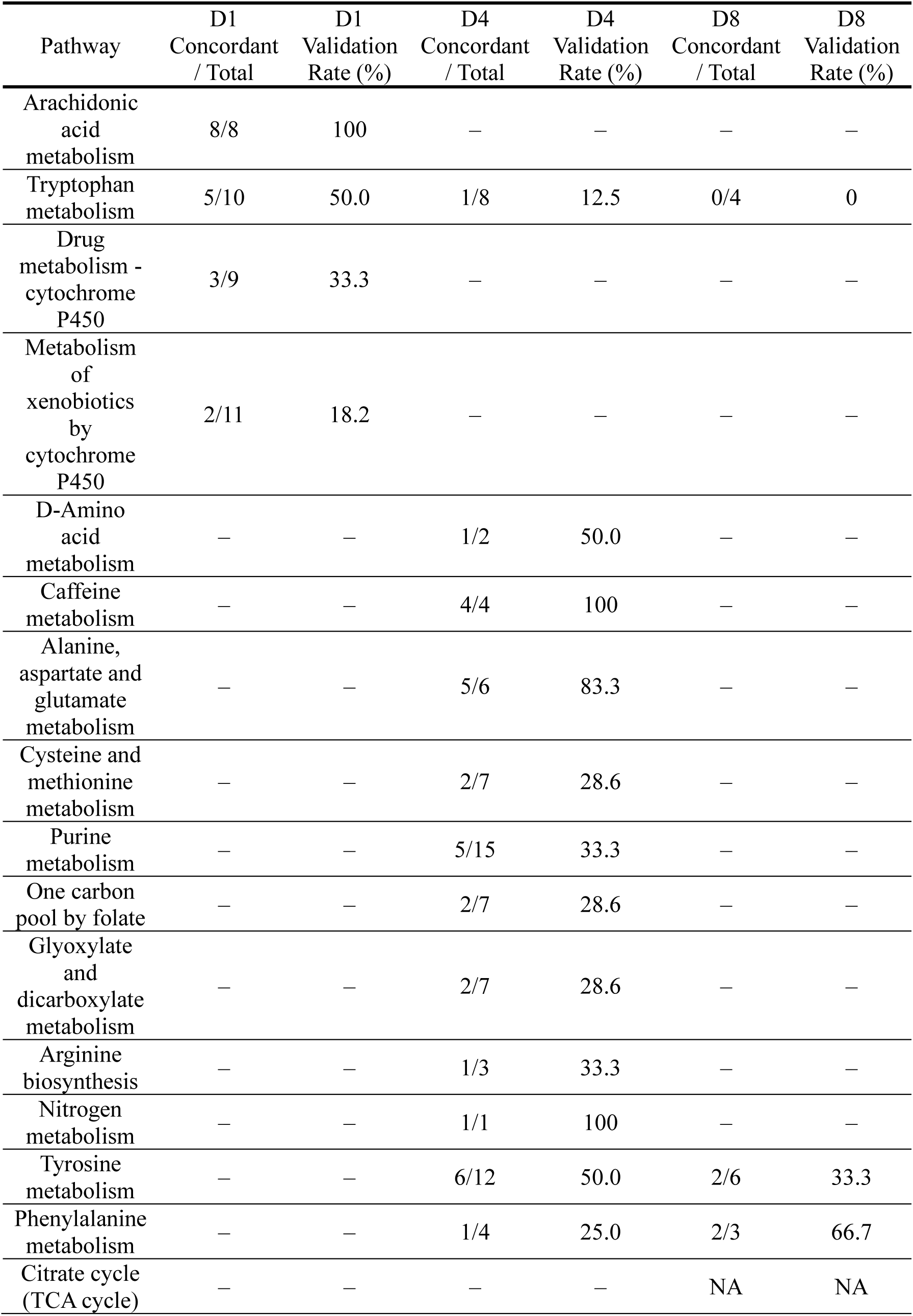
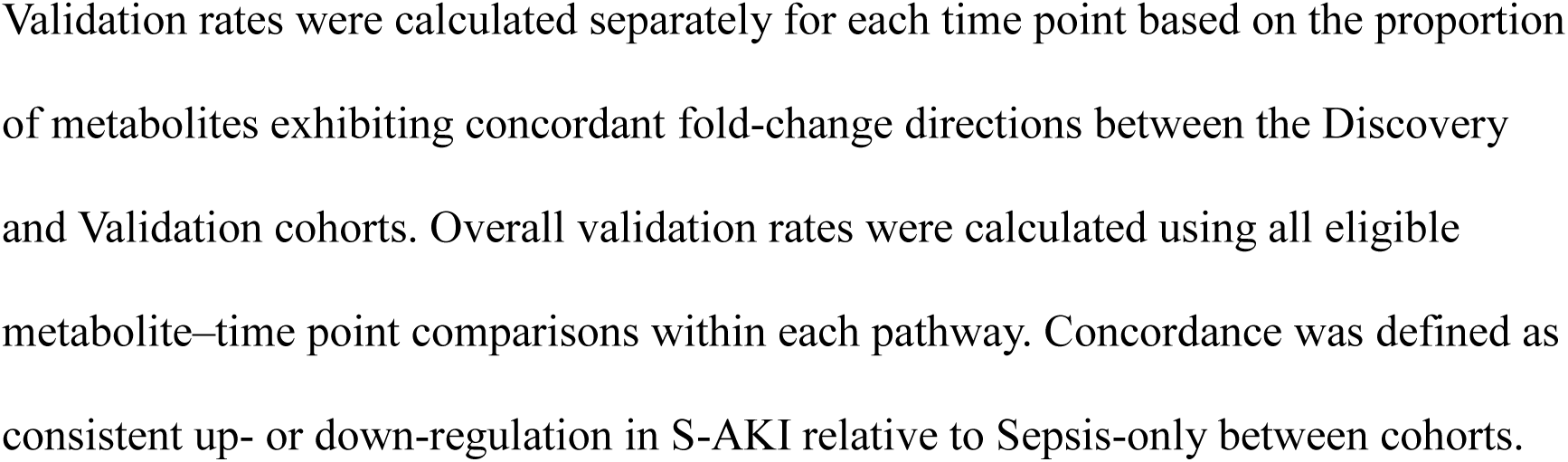
Summary of pathway validation between the Discovery and Validation cohorts of the metabolomic dataset.

| Pathway | D1<br>Concordant<br>/ Total | D1<br>Validation<br>Rate (%) | D4<br>Concordant<br>/ Total | D4<br>Validation<br>Rate (%) | D8<br>Concordant<br>/ Total | D8<br>Validation<br>Rate (%) |
| --- | --- | --- | --- | --- | --- | --- |
| Arachidonic<br>acid<br>metabolism | 8/8 | 100 | – | – | – | – |
| Tryptophan<br>metabolism | 5/10 | 50.0 | 1/8 | 12.5 | 0/4 | 0 |
| Drug<br>metabolism -<br>cytochrome<br>P450 | 3/9 | 33.3 | – | – | – | – |
| Metabolism<br>of<br>xenobiotics<br>by<br>cytochrome<br>P450 | 2/11 | 18.2 | – | – | – | – |
| D-Amino<br>acid<br>metabolism | – | – | 1/2 | 50.0 | – | – |
| Caffeine<br>metabolism | – | – | 4/4 | 100 | – | – |
| Alanine,<br>aspartate and<br>glutamate<br>metabolism | – | – | 5/6 | 83.3 | – | – |
| Cysteine and<br>methionine<br>metabolism | – | – | 2/7 | 28.6 | – | – |
| Purine<br>metabolism | – | – | 5/15 | 33.3 | – | – |
| One carbon<br>pool by folate | – | – | 2/7 | 28.6 | – | – |
| Glyoxylate<br>and<br>dicarboxylate<br>metabolism | – | – | 2/7 | 28.6 | – | – |
| Arginine<br>biosynthesis | – | – | 1/3 | 33.3 | – | – |
| Nitrogen<br>metabolism | – | – | 1/1 | 100 | – | – |
| Tyrosine<br>metabolism | – | – | 6/12 | 50.0 | 2/6 | 33.3 |
| Phenylalanine<br>metabolism | – | – | 1/4 | 25.0 | 2/3 | 66.7 |
| Citrate cycle<br>(TCA cycle) | – | – | – | – | NA | NA |

Validation rates were calculated separately for each time point based on the proportion of metabolites exhibiting concordant fold-change directions between the Discovery and Validation cohorts. Overall validation rates were calculated using all eligible metabolite–time point comparisons within each pathway. Concordance was defined as consistent up- or down-regulation in S-AKI relative to Sepsis-only between cohorts.

## Discussion

The novelty of our study lies in being the first longitudinal investigation to apply comprehensive, non-targeted, bottom-up multi-omics analysis of human urinary extracellular vesicles (uEVs) to characterize the temporal molecular progression of S-AKI. Unlike previous studies that primarily compared AKI and non-AKI groups at a single time point[26, 27], our longitudinal design enabled dynamic tracking of molecular pathway alterations throughout disease progression, revealing stage-specific biological processes from early inflammatory responses to subsequent metabolic reprogramming and late-stage bioenergetic dysfunction.

Urinary EVs have emerged as valuable analytical tools in nephrology because they carry molecular cargo derived directly from renal cells and reflect ongoing pathophysiological changes within the kidney. Previous investigations have identified several urinary EV biomarkers for AKI, including CD35, Fetuin-A, complement proteins, and activating transcription factor 3 (ATF3), supporting the diagnostic potential of uEVs in kidney injury [26–29]. However, these studies focused primarily on individual biomarkers or single-omics approaches. In contrast, our study combines longitudinal proteomics and metabolomics with pathway-level validation, enabling a systems-level characterization of the molecular events underlying S-AKI progression[19]. This integrative strategy provides both mechanistic insight and a framework for identifying biologically relevant pathways and candidate biomarkers across different stages of disease progression.

The validated proteomic and metabolomic pathways and integrated interpretations were summarized in **Figure 5**. Collectively, these pathways revealed a temporal transition from early inflammatory activation to subsequent metabolic adaptation during S-AKI progression. At D1, proteomic analysis identified enrichment of complement and coagulation cascades, ferroptosis, HIF-1 signaling, and sphingolipid metabolism, whereas metabolomic analysis validated arachidonic acid metabolism and tryptophan metabolism. Together, these pathways indicate that the earliest molecular response to S-AKI is characterized by activation of innate immunity, coagulation, inflammatory lipid signaling, and immune-associated metabolic remodeling.

**Figure 5.**
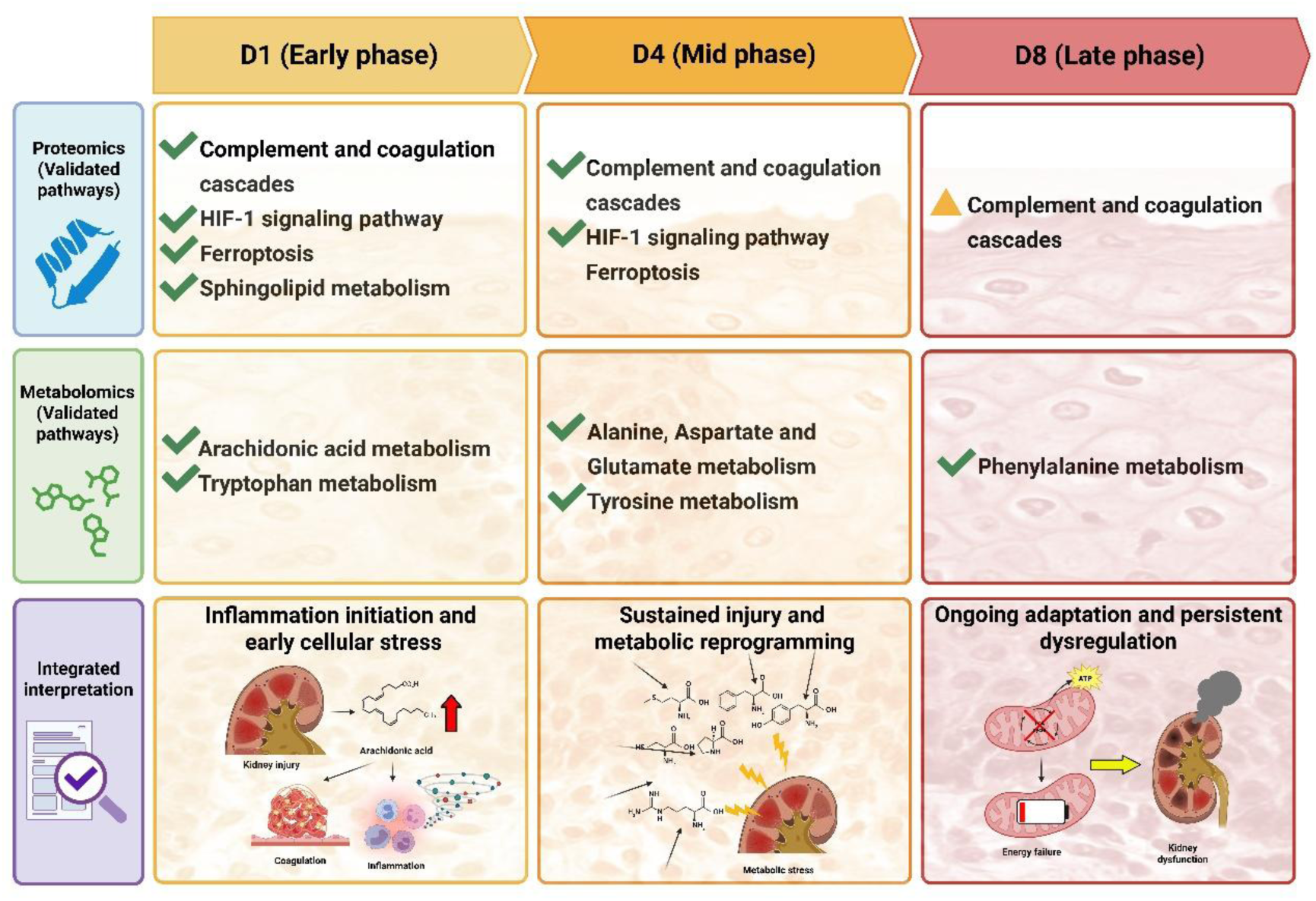
Longitudinal progression of validated proteomic and metabolomic pathways in S-AKI. This figure summarizes the temporal progression of validated proteomic and metabolomic pathways during S-AKI. In the early phase (D1), inflammatory, coagulation, hypoxia, ferroptotic, and lipid metabolic pathways are activated, accompanied by alterations in arachidonic acid and tryptophan metabolism. In the mid-phase (D4), inflammatory signaling persists while amino acid metabolism becomes the predominant metabolic feature, indicating metabolic reprogramming. By the late phase (D8), phenylalanine metabolism remains the only validated pathway, suggesting persistent metabolic dysregulation during disease progression. Pathways that met the predefined validation criteria are indicated by a check mark (✓), whereas pathways with marginal concordance rates are indicated by a triangle (▴).

Complement activation and coagulation dysfunction are well-recognized contributors to endothelial injury, microvascular thrombosis, tubular epithelial damage, and organ dysfunction during sepsis[30], while activation of the HIF-1 signaling pathway reflects cellular adaptation to tissue hypoxia and inflammatory stress[31]. In parallel, ferroptosis has emerged as an important mechanism of tubular cell death in experimental and clinical AKI[32], and sphingolipid metabolism has been implicated in inflammatory signaling and apoptosis in kidney injury [33]. Consistent with these proteomic findings, arachidonic acid-derived prostaglandins and leukotrienes amplify inflammatory responses, leukocyte recruitment, and vascular dysfunction, whereas dysregulated tryptophan metabolism likely reflects inflammation-associated immune-metabolic adaptation through altered kynurenine and serotonin metabolism [34, 35].

During the mid-phase (D4), the validated pathways shifted toward metabolic reprogramming. Proteomic validation of HIF-1 signaling and ferroptosis remained highly consistent, suggesting persistent hypoxia, oxidative stress, and iron-dependent lipid peroxidation. Concurrently, metabolomic analysis demonstrated reproducible alterations in alanine, aspartate and glutamate metabolism, tyrosine metabolism, and caffeine metabolism. Among these, alanine, aspartate and glutamate metabolism showed the highest concordance, supporting glutamate-centered amino acid remodeling as a major metabolic feature during established S-AKI. Glutamate serves as a metabolic hub connecting amino acid transamination, glutathione synthesis, nitrogen metabolism, and replenishment of tricarboxylic acid-cycle intermediates, suggesting increased metabolic demand under sustained inflammatory stress [36]. Alterations in tyrosine metabolism further suggest disturbed aromatic amino acid and catecholamine metabolism during disease progression[37, 38], whereas caffeine metabolism, despite complete validation, was interpreted cautiously because it is strongly influenced by dietary intake rather than representing a disease-specific mechanism.

By D8, no proteomic pathway satisfied the predefined validation criterion, and phenylalanine metabolism was the only validated metabolic pathway. Since disturbances of phenylalanine metabolism have previously been associated with impaired hepatic and renal metabolism as well as systemic inflammation [37, 38], our findings may indicate persistent metabolic dysregulation during late-stage S-AKI. However, because the validation cohort at D8 was markedly limited and unbalanced, these late-stage metabolic findings should be regarded as exploratory. Overall, the validated pathways support a temporal model in which early inflammatory and immune responses gradually transition into extensive amino acid metabolic remodeling during S-AKI progression.

This study has several limitations. First, the relatively small sample size and single-center recruitment may limit the generalizability and external validity of the findings and reduce the statistical power to detect subtle proteomic and metabolomic differences. Second, although urinary EV proteomics provides valuable information for biomarker discovery and pathway interpretation, the limited number of differentially expressed proteins restricts comprehensive pathway analysis and does not establish causal relationships between the identified molecular alterations and S-AKI progression. Third, metabolites were identified using a non-targeted metabolomics workflow and annotated at MSI level-3 confidence[25]. Future studies incorporating authentic reference standards and confirmatory MS/MS analyses will further improve metabolite identification confidence. Moreover, the use of multiple adducts to maximize metabolite annotation coverage increased the likelihood that a single LC–MS feature would be assigned to multiple putative metabolite identities, introducing annotation ambiguity and occasionally leading to heterogeneous validation directions. Pathway validation was based primarily on concordance of fold-change direction between the discovery and validation cohorts rather than statistical significance alone. This limitation was particularly relevant at D8, where the validation cohort was markedly small and unbalanced, with six S-AKI patients and only two sepsis-only patients. The limited sample size substantially reduced statistical power and increased sensitivity to individual observations. Consequently, D8 metabolic findings were considered exploratory and were not used as the principal basis for the mechanistic storyline. Larger multicenter cohorts, targeted metabolite confirmation, and functional studies are required to verify these pathway alterations.

## Conclusion

This study represents the first longitudinal multi-omics investigation of human urinary extracellular vesicles to characterize the temporal molecular progression of S-AKI. By integrating proteomic and metabolomic analyses with an independent validation cohort, we identified reproducible pathway alterations associated with distinct stages of disease progression. Early S-AKI was characterized by inflammatory and immune-metabolic activation, whereas the mid-phase was dominated by glutamate-centered amino acid metabolic reprogramming. These findings provide a temporal molecular framework for S-AKI pathogenesis and demonstrate the potential of urinary EV-based multi-omics for biomarker discovery and mechanistic investigation, thereby facilitating future precision diagnostic and therapeutic strategies.

## List of abbreviations

AKI: Acute kidney injury
AUC: Area under the curve
BUN: Blood urea nitrogen
CKD: Chronic kidney disease
DAMP: Damage-associated molecular patterns
DEP: Differentially expressed proteins
DEF: Differentially expressed features
EV: Extracellular vesicles
FC: Fold change
GO: Gene ontology
ICU: Intensive care unit
KDIGO: Kidney disease: Improving global outcomes
KEGG: Kyoto Encyclopedia of Genes and Genomes
MVE: Multivesicular endosomes
NAPSA: Napsin-A aspartic
NTA: Nano-particle tracking analysis
PAMP: Pathogen-associated molecular patterns
PRR: Pattern recognition receptors
ROC: Receiver operating characteristic
ROS: Reactive oxygen species
SOFA: Sequential organ failure assessment
SPE: Solid-phase extraction
TEC: Tubular epithelial cells
TEM: Transmission electron microscopy
TLR: Toll-like receptors

## Declarations

## Ethics approval and consent to participate

This study was approved by the Joint Institutional Review process of the Taipei Medical University – Joint Institutional Review Board No. N202404097). Signed informed consent was obtained from all participants recruited in the study.

## Consent for publication

Not applicable

## Availability of data and materials

All data related to this article are disclosed or available upon request from the corresponding author.

## Authors’ contributions

Conceptualization: TYC, ILT, CCK; Investigation: TYC, GYC, TIW, SYW; Formal analysis: TYC, YCS, CYC. Visualization: ILT, LYH, IJC; Validation: YCL, HHC, WCC; Resources: GYC, TIW, SYW; Writing—Original Draft: TYC, ILT, CCK; Writing—Review & Editing: HHC, WCC; Supervision: MSW, CCK; Project administration: ILT, CCK; Funding acquisition: ILT, CCK.

## Date statement

Most of the data and information supporting the findings of this study are available within the main text. The raw mass spectrometry proteomics data related to Figures 2–7 have been deposited in the ProteomeXchange Consortium via the PRIDE partner repository under the dataset identifier PXD063789. Additional datasets are available from the corresponding author, CK, on reasonable request.

## Acknowledgments

We thank the mass spectrometry technical research services provided by Consortia of Key Technologies and Instrumentation Center, National Taiwan University.

## References

1. Lin, Y.-H., et al., Global proteome and phosphoproteome characterization of sepsis-induced kidney injury. Molecular & Cellular Proteomics, 2020. 19(12): p. 2030–2047.

2. Albert, C., et al., Neutrophil gelatinase-associated lipocalin measured on clinical laboratory platforms for the prediction of acute kidney injury and the associated need for dialysis therapy: a systematic review and meta-analysis. American Journal of Kidney Diseases, 2020. 76(6): p. 826–841. e1.

3. Haase, M., et al., Accuracy of neutrophil gelatinase-associated lipocalin (NGAL) in diagnosis and prognosis in acute kidney injury: a systematic review and meta-analysis. American journal of kidney diseases, 2009. 54(6): p. 1012–1024.

4. Geng, J., et al., The value of kidney injury molecule 1 in predicting acute kidney injury in adult patients: a systematic review and Bayesian meta-analysis. Journal of Translational Medicine, 2021. 19(1): p. 105.

5. Shao, X., et al., Diagnostic value of urinary kidney injury molecule 1 for acute kidney injury: a meta-analysis. PloS one, 2014. 9(1): p. e84131.

6. Liu, C., et al., The diagnostic accuracy of urinary [TIMP-2]·[IGFBP7] for acute kidney injury in adults: A PRISMA-compliant meta-analysis. Medicine, 2017. 96(27): p. e7484.

7. Vijayan, A., et al., Clinical use of the urine biomarker [TIMP-2]×[IGFBP7] for acute kidney injury risk assessment. American Journal of Kidney Diseases, 2016. 68(1): p. 19–28.

8. Khorashadi, M., et al., Proenkephalin: a new biomarker for glomerular filtration rate and acute kidney injury. Nephron, 2020. 144(12): p. 655–661.

9. Caironi, P., et al., Circulating proenkephalin, acute kidney injury, and its improvement in patients with severe sepsis or shock. Clinical chemistry, 2018. 64(9): p. 1361–1369.

10. Zdziechowska, M., et al., Serum NGAL, KIM-1, IL-18, L-FABP: new biomarkers in the diagnostics of acute kidney injury (AKI) following invasive cardiology procedures. International urology and nephrology, 2020. 52(11): p. 2135–2143.

11. Susantitaphong, P., et al., Performance of urinary liver-type fatty acid–binding protein in acute kidney injury: a meta-analysis. American Journal of Kidney Diseases, 2013. 61(3): p. 430–439.

12. Khawaja, S., et al., The utility of neutrophil gelatinase-associated Lipocalin (NGAL) as a marker of acute kidney injury (AKI) in critically ill patients. Biomarker research, 2019. 7: p. 1–6.

13. Xu, X., et al., Epidemiology and clinical correlates of AKI in Chinese hospitalized adults. Clinical Journal of the American Society of Nephrology, 2015. 10(9): p. 1510–1518.

14. Peerapornratana, S., et al., Acute kidney injury from sepsis: current concepts, epidemiology, pathophysiology, prevention and treatment. Kidney international, 2019. 96(5): p. 1083–1099.

15. Van Niel, G., G. d’Angelo, and G. Raposo, Shedding light on the cell biology of extracellular vesicles. Nature reviews Molecular cell biology, 2018. 19(4): p. 213–228.

16. Chen, Y.-F., et al., Exosomes: A review of biologic function, diagnostic and targeted therapy applications, and clinical trials. Journal of biomedical science, 2024. 31(1): p. 67.

17. Kalluri, R. and V.S. LeBleu, The biology, function, and biomedical applications of exosomes. Science, 2020. 367(6478): p. eaau6977.

18. Merchant, M.L., et al., Isolation and characterization of urinary extracellular vesicles: implications for biomarker discovery. Nature Reviews Nephrology, 2017. 13(12): p. 731–749.

19. Qiao, J. and L. Cui, Multi-omics techniques make it possible to analyze sepsis-associated acute kidney injury comprehensively. Frontiers in Immunology, 2022. 13: p. 905601.

20. Kellum, J.A., et al., Kidney disease: improving global outcomes (KDIGO) acute kidney injury work group. KDIGO clinical practice guideline for acute kidney injury. Kidney international supplements, 2012. 2(1): p. 1–138.

21. Biosciences, S. SmartSEC® HT EV Isolation System for Serum & Plasma User Manual. 2019 [cited 2024 February 26, 2024]; Available from: https://www.systembio.com/wp/wp-content/uploads/2020/10/MANUAL_SmartSEC_HT_EV_Isolation_System_Plasma-Serum.pdf.

22. Kowal, E.J., et al., Extracellular vesicle isolation and analysis by western blotting. Extracellular vesicles: methods and protocols, 2017: p. 143–152.

23. Cheng, Y.-H., et al., Multiplexed antibody glycosylation profiling using dual enzyme digestion and liquid chromatography-triple quadrupole mass spectrometry method. Molecular & Cellular Proteomics, 2024. 23(2): p. 100710.

24. Varghese, R.S., et al., Ion annotation-assisted analysis of LC-MS based metabolomic experiment. Proteome science, 2012. 10(Suppl 1): p. S8.

25. Schrimpe-Rutledge, A.C., et al., Untargeted metabolomics strategies—challenges and emerging directions. Journal of the American Society for Mass Spectrometry, 2016. 27(12): p. 1897–1905.

26. Panich, T., et al., Urinary exosomal activating transcriptional factor 3 as the early diagnostic biomarker for sepsis-induced acute kidney injury. BMC nephrology, 2017. 18: p. 1–13.

27. Li, N., et al., Single urinary extracellular vesicle proteomics identifies complement receptor CD35 as a biomarker for sepsis-associated acute kidney injury. Nature Communications, 2025. 16(1): p. 6960.

28. Zhou, H., et al., Exosomal Fetuin-A identified by proteomics: a novel urinary biomarker for detecting acute kidney injury. Kidney international, 2006. 70(10): p. 1847–1857.

29. Awdishu, L., et al., Urinary exosomes identify inflammatory pathways in vancomycin associated acute kidney injury. International journal of molecular sciences, 2021. 22(6): p. 2784.

30. Wei, L., et al. Role of the Complement System in Acute Kidney Injury: A Narrative Review. in Mayo Clinic Proceedings. 2025. Elsevier.

31. Li, W., et al., Mitochondria bridge HIF signaling and ferroptosis blockage in acute kidney injury. Cell death & disease, 2022. 13(4): p. 308.

32. Zhang, J., et al., The role of ferroptosis in acute kidney injury. Frontiers in Molecular Biosciences, 2022. 9: p. 951275.

33. Dupre, T.V. and L.J. Siskind, The role of sphingolipids in acute kidney injury. Advances in biological regulation, 2018. 70: p. 31–39.

34. Wang, T., et al., Arachidonic acid metabolism and kidney inflammation. International journal of molecular sciences, 2019. 20(15): p. 3683.

35. Krupa, A., M.M. Krupa, and K. Pawlak, Kynurenine pathway—an underestimated factor modulating innate immunity in sepsis-induced acute kidney injury? Cells, 2022. 11(16): p. 2604.

36. Turk, H., E. Temiz, and I. Koyuncu, Metabolic reprogramming in sepsis-associated acute kidney injury: insights from lipopolysaccharide-induced oxidative stress and amino acid dysregulation. Molecular Biology Reports, 2025. 52(1): p. 52.

37. Xu, W., et al., Integrating metabolomics and machine learning with in silico analysis to identify early biomarkers and molecular interactions in sepsis-associated acute kidney injury. Scientific Reports, 2026.

38. Qu, X., et al., Identification of key metabolites during cisplatin-induced acute kidney injury using an HPLC-TOF/MS-based non-targeted urine and kidney metabolomics approach in rats. Toxicology, 2020. 431: p. 152366.

