## Supplementary material for "Integrated Proteo-Metabolomics of Urinary Extracellular Vesicles Reveals Early Molecular Divergence and Temporal Pathogenesis of Sepsis-Associated AKI": Table S1

**Table S4. Validation of pathway-enriched proteins identified in the Discovery cohort using an independent Validation cohort.**

|  | **Discovery cohort** | | | | | | **Validation cohort** | | | | | | **Validation status ^d^** | | |
| --- | --- | --- | --- | --- | --- | --- | --- | --- | --- | --- | --- | --- | --- | --- | --- |
|  | **D1** | | **D4** | | **D8** | | **D1** | | **D4** | | **D8 ^c^** | | **D1** | **D4** | **D8** |
| Protein | p value ^a^ | FC ^b^ | p value | FC | p value | FC | p value | FC | p value | FC | p value | FC |  |  |  |
| **Complement and coagulation cascades** |  | | | | | | | | | | | |  | | |
| CFD | 0.046 | 16.56 | 0.023 | 69.76 | 0.002 | 87.13 | 0.318 | 0.54 | 0.682 | 0.73 | 0.889 | 16.34 |  |  |  |
| SERPINA5 | 0.001 | 0.11 | 0.005 | 0.11 | 0.088 | 0.35 | 0.560 | 4.98 | 0.537 | 0.59 | 0.667 | 0.13 |  |  |  |
| A2M | 0.024 | 3.48 | 0.028 | 2.87 | - ^e^ | - | 0.138 | 2.25 | 0.964 | 0.97 | - | - |  |  | - |
| C3 | 0.074 | 2.16 | 0.004 | 2.165 | - | - | 0.285 | 1.48 | 0.163 | 2.16 | - | - |  |  | - |
| C7 | 0.038 | 4.31 | 0.005 | 8.94 | - | - | 0.154 | 1.96 | 0.249 | 2.52 | - | - |  |  | - |
| C9 | 0.048 | 2.02 | - | - | 0.013 | 7.03 | 0.361 | 1.05 | - | - | >0.999 | 0.96 |  | - |  |
| CFB | 0.080 | 0.61 | 0.002 | 5.9 | - | - | 0.011 | 2.09 | 0.072 | 2.44 | - | - |  |  | - |
| F2 | 0.041 | 2.04 | - | - | 0.049 | 3.76 | 0.043 | 2.14 | - | - | 0.278 | 0.34 |  | - |  |
| C8G | - | - | 0.009 | 2.47 | 0.027 | 8.76 | - | - | 0.36285 | 1.74 | >0.999 | N/A | - |  |  |
| SERPINA1 | 0.090 | 1.74 | - | - | - | - | 0.533 | 1.42 | - | - | - | - |  | - | - |
| PROS1 | - | - | 0.047 | 1.60 | - | - | - | - | 0.217 | 35.26 | - | - | - |  | - |
| CFI | - | - | 0.025 | 6.82 | - | - | - | - | 0.178 | 2.85 | - | - | - |  | - |
| FGG | - | - | - | - | 0.088 | 14.27 | - | - | - | - | 0.889 | 17.61 | - | - |  |
| VTN | - | - | - | - | 0.076 | 3.44 | - | - | - | - | >0.999 | 0.80 | - | - |  |
| **Sphingolipid metabolism** |  | | | | | | | | | | | |  |  |  |
| ASAH1 | 0.033 | 4.79 | - | - | - | - | 0.020 | 25.31 | - | - | - | - |  | - | - |
| GLB1 | 0.079 | 0.33 | - | - | - | - | 0.478 | 35.36 | - | - | - | - |  | - | - |
| PSAP | 0.034 | 4.79 | - | - | - | - | 0.400 | 2.23 | - | - | - | - |  | - | - |
| GM2A | 0.002 | 0.16 | - | - | - | - | N/A | N/A | - | - | - | - |  | - | - |
| **HIF-1 signaling pathway** |  | | | | | | | | | | | |  |  |  |
| EGF | 0.007 | 0.27 | 0.071 | 0.13 | - | - | 0.933 | 0.22 | 0.717 | 0.68 | - | - |  |  | - |
| LDHA | 0.031 | 2.58 | 0.053 | 2.90 | - | - | 0.941 | 6.09 | 0.212 | 8.71 | - | - |  |  | - |
| TF | 0.098 | 1.64 | 0.045 | 2.55 | - | - | 0.226 | 0.91 | 0.173 | 2.34 | - | - |  |  | - |
| **Ferroptosis** |  | | | | | | | | | | | |  |  |  |
| CP | 0.022 | 2.73 | 0.019 | 2.90 | - | - | 0.112 | 1.35 | 0.230 | 1.97 | - | - |  |  | - |
| TF | 0.098 | 1.64 | 0.045 | 2.55 | - | - | 0.226 | 0.91 | 0.173 | 2.34 | - | - |  |  | - |
| SLC3A2 | 0.014 | 0.34 | - | - | - | - | 0.004 | 0.04 | - | - | - | - |  | - | - |

^a^ P values were calculated using the Wilcoxon rank-sum test for comparisons between the S-AKI and Sepsis-only groups.

^b^ Fold change (FC) was calculated as the ratio of mean protein abundance in the S-AKI group relative to the Sepsis-only group.

^c^ Due to insufficient sample size in the Validation cohort at D8 (7 S-AKI and 2 Sepsis-only samples), Mann–Whitney U tests were performed instead of Wilcoxon rank-sum tests.

^d^ Proteins were considered concordantly validated when the direction of fold change (up- or down-regulation in S-AKI relative to Sepsis-only) was consistent between the Discovery and Validation cohorts at the corresponding time point. Concordant and discordant validation results are highlighted in green and red, respectively. Proteins without corresponding fold-change data in the validation cohort are highlighted in gray and treated as discordant for the calculation of the concordance rate.

^e^ – indicates that the protein was not identified as a pathway-enriched candidate (p < 0.1 and FC > 1.3) at the corresponding time point in the Discovery cohort and was therefore not included in the validation analysis.

AKI, acute kidney injury; FC, fold change; S-AKI, sepsis-associated acute kidney injury; HIF, hypoxia-inducible factor; N/A, not available.
