## Supplementary material for "Integrated Proteo-Metabolomics of Urinary Extracellular Vesicles Reveals Early Molecular Divergence and Temporal Pathogenesis of Sepsis-Associated AKI": Table S2

**Table S5. Validation of pathway-enriched metabolites identified in the Discovery cohort using an independent Validation cohort.**

|  | **Discovery cohort** | | | | | | **Validation cohort** | | | | | | **Validation status ^d^** | | |
| --- | --- | --- | --- | --- | --- | --- | --- | --- | --- | --- | --- | --- | --- | --- | --- |
|  | **D1** | | **D4** | | **D8** | | **D1** | | **D4** | | **D8 ^c^** | | **D1** | **D4** | **D8** |
| Metaabolites | p value ^a^ | FC ^b^ | p value | FC | p value | FC | p value | FC | p value | FC | p value | FC |  |  |  |
| **Arachidonic acid metabolism** |  | | | | | | | | | | | |  | | |
| Prostaglandin H2 | 0.044 | 1.4 | - | - | - | - | 0.284 | 1.28 | - | - | - | - |  | - | - |
| Prostaglandin D2 | 0.044 | 1.4 | - | - | - | - | 0.284 | 1.28 | - | - | - | - |  | - | - |
| Prostaglandin E2 | 0.044 | 1.4 | - | - | - | - | 0.284 | 1.28 | - | - | - | - |  | - | - |
| 20-OH-Leukotriene B4 | 0.044 | 1.4 | - | - | - | - | 0.284 | 1.28 | - | - | - | - |  | - | - |
| Prostaglandin I2 | 0.044 | 1.4 | - | - | - | - | 0.284 | 1.28 | - | - | - | - |  | - | - |
| Thromboxane A2 | 0.044 | 1.4 | - | - | - | - | 0.284 | 1.28 | - | - | - | - |  | - | - |
| 15-Keto-prostaglandin F2alpha | 0.044 | 1.4 | - | - | - | - | 0.284 | 1.28 | - | - | - | - |  | - | - |
| (5Z)-11alpha-Hydroxy-9,15-dioxoprostanoate | 0.044 | 1.4 | - | - | - | - | 0.284 | 1.28 | - | - | - | - |  | - | - |
| **Tryptophan metabolism** |  | | | | | | | | | | | | | | |
| L-Tryptophan | 0.028 | 0.55 | 0.048 | 2.25 | - | - | NA | NA | 0.382 | 1.27 | - | - |  |  | - |
| 5-Hydroxyindoleacetate | 0.029 | 0.54 | 0.011 | 0.07 | 0.034 | 0.2 | 0.213 | 0.65 | 0.250, 0.448, 0.884 | 0.61, 16.12, 0.93 | 0.929 | 9.3 |  |  |  |
| 2-Oxoadipate | - | - | - | - | 0.005 | 0.51 | - | - | - | - | NA | NA | - | - |  |
| 5-Hydroxy-L-tryptophan | 0.013 | 0.31 | 0.041 | 0.18 | - | - | 0.698 | 0.11 | 0.438 | 151.17 | - | - |  |  | - |
| 3-Hydroxyanthranilate | 0.001 | 1.93 | 0.033 | 1.7 | 0.013 | 0.7 | 0.097 | 2.05 | NA | NA | 0.286 | 7.61 |  |  |  |
| L-Formylkynurenine | - | - | 0.039; 0.044; 0.047 | 7.78; 0.02; 0.08 | - | - | - | - | 0.151 | 0.49 | - | - | - |  | - |
| 6-Hydroxymelatonin | - | - | 0.011; 0.023 | 0.11; 0.22 | 0.034 | 1.76 | - | - | NA | NA | 0.286 | 0.14 | - |  |  |
| L-Kynurenine | 0.029 | 0.54 | - | - | - | - | NA | NA | - | - | - | - |  | - | - |
| 2-Amino-3-carboxymuconate semialdehyde | - | - | 0.012 | 0.03 | - | - | - | - | NA | NA | - | - | - |  | - |
| Indolepyruvate | 0.0005 | 0.28 | - | - | - | - | NA | NA | - | - | - | - |  | - | - |
| Formyl-5-hydroxykynurenamine | 0.029 | 0.54 | - | - | - | - | NA | NA | - | - | - | - |  | - | - |
| Formylanthranilate | - | - | 0.048; 0.049 | 0.19; 43.15 | - | - | - | - | 0.205 | 0.007 | - | - | - |  | - |
| 5-Methoxyindoleacetate | 0.006 | 0.74 | - | - | - | - | 0.349, 0.398 | 1.82, 1.86 | - | - | - | - |  | - | - |
| Formyl-N-acetyl-5-methoxykynurenamine | - | - | 0.01 | 0.003 | - | - | - | - | NA | NA | - | - | - |  | - |
| Anthranilate | 0.027 | 1.52 | 0.014 | 2.03 | - | - | 0.468 | 29.99 | 0.271, 0.918 | 0.40, 0.94 | - | - |  |  | - |
| 4-(2-Amino-3-hydroxyphenyl)-2,4-dioxobutanoate | 0.021 | 0.41 | - | - | - | - | 0.067, 0.484 | 0.41, 0.82 | - | - | - | - |  | - | - |
| N-Methyltryptamine | - | - | 0.007 | 0.44 | - | - | - | - | NA | NA | - | - | - |  | - |
| **Drug metabolism - cytochrome P450** |  | | | | | | | | | | | | | | |
| Citalopram aldehyde | 0.005 | 0.69 | - | - | - | - | NA | NA | - | - | - | - |  | - | - |
| 2-Phenyl-1,3-propanediol monocarbamate | 0.042 | 0.71 | - | - | - | - | 0.924, 0.384, 0.200, 0.536 | 1.02, 1.48, 1.72, 0.83 | - | - | - | - |  | - | - |
| 4-Hydroxy-5-phenyltetrahydro-1,3-oxazin-2-one | 0.024 | 0.53 | - | - | - | - | 0.796 | 0.94 | - | - | - | - |  | - | - |
| 3-Carbamoyl-2-phenylpropionaldehyde | 0.024 | 0.53 | - | - | - | - | 0.796 | 0.94 | - | - | - | - |  | - | - |
| 3-Hydroxylidocaine | 0.032 | 0.65 | - | - | - | - | 0.36 | 1.42 | - | - | - | - |  | - | - |
| 4-Ketocyclophosphamide | 0.014 | 0.4 | - | - | - | - | NA | NA | - | - | - | - |  | - | - |
| Alcophosphamide | 0.032 | 0.52 | - | - | - | - | 0.476 | 0.66 | - | - | - | - |  | - | - |
| Morphine-3-glucuronide | 0.008 | 1.35 | - | - | - | - | NA | NA | - | - | - | - |  | - | - |
| 5-Phenyl-1,3-oxazinane-2,4-dione | 0.029 | 0.54 | - | - | - | - | 0.662, 0.923, 0.213 | 1.72, 1.06, 0.65 | - | - | - | - |  | - | - |
| Norcodeine | 0.007 | 0.77 | - | - | - | - | 0.294 | 1.12 | - | - | - | - |  | - | - |
| Morphine-6-glucuronide | 0.008 | 1.35 | - | - | - | - | NA | NA | - | - | - | - |  | - | - |
| **Metabolism of xenobiotics by cytochrome P450** |  | | | | | | | | | | | | | | |
| 4-Bromophenol | 0.032 | 0.61 | - | - | - | - | 0.705, 0.977, 0.887 | 1.02, 1.80, 2.18 | - | - | - | - |  | - | - |
| Bromobenzene-2,3-oxide | 0.032 | 0.61 | - | - | - | - | 0.705, 0.977, 0.887 | 1.02, 1.80, 2.18 | - | - | - | - |  | - | - |
| 2-(S-Glutathionyl)acetyl chloride | 0.006 | 0.7 | - | - | - | - | NA | NA | - | - | - | - |  | - | - |
| Bromobenzene-3,4-oxide | 0.032 | 0.61 | - | - | - | - | 0.705, 0.977, 0.887 | 1.02, 1.80, 2.18 | - | - | - | - |  | - | - |
| Aflatoxin B1-exo-8,9-epoxide | 0.031 | 0.62 | - | - | - | - | 0.796 | 0.94 | - | - | - | - |  | - | - |
| Aflatoxin M1 | 0.031 | 0.62 | - | - | - | - | 0.333 | 11.81 | - | - | - | - |  | - | - |
| Naphthalene | 0.026 | 0.49 | - | - | - | - | 0.18 | 0.12 | - | - | - | - |  | - | - |
| S-(1,2-Dichlorovinyl)glutathione | 0.017; 0.004 | 0.36; 0.50 | - | - | - | - | NA | NA | - | - | - | - |  | - | - |
| S-(2-Chloroacetyl)glutathione | 0.006 | 0.7 | - | - | - | - | NA | NA | - | - | - | - |  | - | - |
| Aflatoxin Q1 | 0.031 | 0.62 | - | - | - | - | 0.333 | 11.81 | - | - | - | - |  | - | - |
| 7-Hydroxymethyl-12-methylbenz[a]anthracene sulfate | 0.025 | 0.68 | - | - | - | - | 0.473, 0.366 | 5.12, 10.00 | - | - | - | - |  | - | - |
| **D-Amino acid metabolism** |  | | | | | | | | | | | | | | |
| D-Glutamine | - | - | 0.044; 0.033; 0.023; 0.013; 0.011 | 0.32; 0.006; 0.007; 0.03; 1.86 | - | - | - | - | 0.485, 0.081 | 0.73, 1.60 | - | - | - |  | - |
| D-Glutamate | - | - | 0.033 | 1.63 | - | - | - | - | 0.414, 0.363 | 229.3, 4.40 | - | - | - |  | - |
| D-Arginine | - | - | 0.042 | 0.13 | - | - | - | - | 0.295 | 1.32 | - | - | - |  | - |
| D-Proline | - | - | 0.048 | 0.6 | - | - | - | - | 0.758, 0.670, 0.335, 0.514, 0.823, 0.336, 0.501, 0.387, 0.163, 0.317, 0.432 | 0.87, 0.69, 1.47, 1.33, 1.08, 0.60, 0.78, 2.26, 0.64, 0.68, 0.69 | - | - | - |  | - |
| cis-4-Hydroxy-D-proline | - | - | 0.037 | 1.9 | - | - | - | - | 0.368, 0.932, 0.228 | 0.67, 0.94, 1.55 | - | - | - |  | - |
| 5-Amino-2-oxopentanoic acid | - | - | 0.037 | 1.9 | - | - | - | - | 0.932, 0.228, 0.368 | 0.94, 1.55, 0.67 | - | - | - |  | - |
| 1-Pyrroline-2-carboxylate | - | - | 0.017 | 0.14 | - | - | - | - | 0.314, 0.168, 0.191 | 1.53, 0.69, 0.64 | - | - | - |  | - |
| 1-Pyrroline-4-hydroxy-2-carboxylate | - | - | 0.026; 0.033 | 1.81; 1.63 | - | - | - | - | 0.568, 0.434, 0.760, 0.791, 0.170, 0.717 | 2.44, 3.31, 1.07, 1.16, 1.53, 0.92 | - | - | - |  | - |
| **Caffeine metabolism** |  | | | | | | | | | | | | | | |
| 1,7-Dimethylxanthine | - | - | 0.046 | 0.13 | - | - | - | - | 0.409, 0.160, 0.459 | 0.46, 0.24, 0.44 | - | - | - |  | - |
| 1-Methylxanthine | - | - | 0.012 | 6.05 | - | - | - | - | 0.245, 0.218, 0.112 | 0.32, 0.20, 4.00 | - | - | - |  | - |
| Theobromine | - | - | 0.046 | 0.13 | - | - | - | - | 0.459, 0.409, 0.160 | 0.44, 0.46, 0.24 | - | - | - |  | - |
| 7-Methylxanthine | - | - | 0.012 | 6.05 | - | - | - | - | 0.112, 0.245, 0.218 | 4.00, 0.32, 0.20 | - | - | - |  | - |
| 3,7-Dimethyluric acid | - | - | 0.020; 0.042 | 0.08; 0.08 | - | - | - | - | 0.419 | 0.49 | - | - | - |  | - |
| 1,7-Dimethyluric acid | - | - | 0.020; 0.042 | 0.08; 0.08 | - | - | - | - | 0.419 | 0.49 | - | - | - |  | - |
| **Alanine, aspartate and glutamate metabolism** |  | | | | | | | | | | | | | | |
| N-Acetyl-L-aspartate | - | - | 0.044 | 2.46 | - | - | - | - | 0.165 | 17.4 | - | - | - |  | - |
| L-Alanine | - | - | 0.012 | 0.12 | - | - | - | - | 0.112, 0.255 | 0.51, 0.74 | - | - | - |  | - |
| Succinate semialdehyde | - | - | 0.036 | 1.31 | - | - | - | - | 0.879, 0.642, 0.147, 0.948, 0.989, 0.694, 0.481 | 1.06, 1.20, 3.07, 0.98, 1.00, 0.93, 0.86 | - | - | - |  | - |
| L-Glutamate | - | - | 0.033 | 1.63 | - | - | - | - | 0.414, 0.363 | 229.3, 4.40 | - | - | - |  | - |
| 4-Aminobutanoate | - | - | 0.003 | 0.02 | - | - | - | - | 0.517, 0.567, 0.883, 0.172, 0.125, 0.307, 0.138, 0.257, 0.168 | 1.10, 0.71, 0.95, 0.34, 0.35, 0.50, 0.59, 0.47, 0.32 | - | - | - |  | - |
| L-Glutamine | - | - | 0.044; 0.033; 0.023; 0.013 0.011 | 0.32; 0.006; 0.007; 0.03; 1.86 | - | - | - | - | 0.081, 0.485 | 1.6, 0.73 | - | - | - |  | - |
| 2-Oxoglutaramate | - | - | 0.013, 0.005 | 1.35, 0.004 | - | - | - | - | 0.081 | 1.6 | - | - | - |  | - |
| (S)-1-Pyrroline-5-carboxylate | - | - | 0.017 | 0.14 | - | - | - | - | 0.168, 0.191, 0.314 | 0.69, 0.64, 1.53 | - | - | - |  | - |
| Citrate | - | - | 0.002; 0.012 | 0.35; 0.11 | - | - | - | - | NA | NA | - | - | - |  | - |
| Pyruvate | - | - | 0.043838 | 0.37 | - | - | - | - | 0.334 | 0.76 | - | - | - |  | - |
| **Cysteine and methionine metabolism** |  | | | | | | | | | | | | | | |
| S-Adenosylmethioninamine | - | - | 0.033 | 0.59 | - | - | - | - | 0.338 | 0.59 | - | - | - |  | - |
| S-Adenosyl-L-methionine | - | - | 0.042 | 1.49 | - | - | - | - | 0.684 | 0.81 | - | - | - |  | - |
| S-Adenosyl-L-homocysteine | - | - | 0.046 | 0.37 | - | - | - | - | NA | NA | - | - | - |  | - |
| 2,3-Diketo-5-methylthiopentyl-1-phosphate | - | - | 0.005 | 2.26 | - | - | - | - | NA | NA | - | - | - |  | - |
| L-Cystine | - | - | 0.028 | 0.7 | - | - | - | - | NA | NA | - | - | - |  | - |
| (S)-2-Aminobutanoate | - | - | 0.003 | 0.02 | - | - | - | - | 0.307, 0.138, 0.257, 0.168, 0.517, 0.567, 0.883, 0.172, 0.125 | 0.50, 0.59, 0.47, 0.32, 1.10, 0.95, 0.34, 0.35 | - | - | - |  | - |
| 2-Oxobutanoate | - | - | 0.036 | 1.31 | - | - | - | - | 0.879, 0.642, 0.147, 0.948, 0.989, 0.694, 0.481 | 1.06, 1.20, 3.07, 0.98, 1.00, 0.93, 0.86 | - | - | - |  | - |
| 3-(Methylthio)propanoate | - | - | 0.011 | 0.61 | - | - | - | - | NA | NA | - | - | - |  | - |
| Dehydroalanine | - | - | 0.033 | 2.21 | - | - | - | - | 0.37, 0.913, 0.144, 0.219 | 0.31, 0.93, 8.77, 12.59 | - | - | - |  | - |
| Pyruvate | - | - | 0.044 | 0.37 | - | - | - | - | 0.334 | 0.76 | - | - | - |  | - |
| **Purine metabolism** |  | | | | | | | | | | | | | | |
| L-Glutamine | - | - | 0.011; 0.013; 0.023; 0.033; 0.044 | 1.86; 0.03; 0.007; 0.006; 0.32 | - | - | - | - | 0.485 | 0.73 | - | - | - |  | - |
| 5'-Phosphoribosylglycinamide | - | - | 0.016 | 3.88 | - | - | - | - | 0.106, 0.353, 0.219, 0.247 | 1.63, 1.30, 1.48, 1.35 | - | - | - |  | - |
| 1-(5'-Phosphoribosyl)-5-amino-4-imidazolecarboxamide | - | - | 0.037 | 0.2 | - | - | - | - | 0.456 | 0.48 | - | - | - |  | - |
| 1-(5-Phospho-D-ribosyl)-5-amino-4-imidazolecarboxylate | - | - | 0.006 | 0.09 | - | - | - | - | NA | NA | - | - | - |  | - |
| AMP | - | - | 0.013 | 0.05 | - | - | - | - | 0.807 | 1.12 | - | - | - |  | - |
| Adenosine | - | - | 0.031 | 0.61 | - | - | - | - | 0.428 | 0.54 | - | - | - |  | - |
| Xanthosine | - | - | 0.038 | 0.63 | - | - | - | - | 0.245 | 1.61 | - | - | - |  | - |
| IDP | - | - | 0.005 | 0.04 | - | - | - | - | NA | NA | - | - | - |  | - |
| GMP | - | - | 0.045 | 0.68 | - | - | - | - | 0.861 | 0.88 | - | - | - |  | - |
| Inosine | - | - | 0.005 | 0.13 | - | - | - | - | NA | NA | - | - | - |  | - |
| Deoxyguanosine | - | - | 0.031 | 0.61 | - | - | - | - | 0.428 | 0.54 | - | - | - |  | - |
| dGMP | - | - | 0.013 | 0.05 | - | - | - | - | 0.807 | 1.12 | - | - | - |  | - |
| Sulfate | - | - | 0.04 | 0.09 | - | - | - | - | NA | NA | - | - | - |  | - |
| Adenylyl sulfate | - | - | 0.023 | 0.14 | - | - | - | - | 0.679, 0.535 | 1.55, 2.16 | - | - | - |  | - |
| Urate | - | - | 0.010; 0.022; 0.032 | 0.15; 0.09; 0.04 | - | - | - | - | NA | NA | - | - | - |  | - |
| 5-Amino-4-imidazolecarboxyamide | - | - | 0.02 | 6.36 | - | - | - | - | NA | NA | - | - | - |  | - |
| **One carbon pool by folate** |  | | | | | | | | | | | | | | |
| Dihydrofolate | - | - | 0.045 | 2.78 | - | - | - | - | NA | NA | - | - | - |  | - |
| 5-Formiminotetrahydrofolate | - | - | 0.012 | 2.5 | - | - | - | - | NA | NA | - | - | - |  | - |
| S-Adenosyl-L-methionine | - | - | 0.042 | 1.49 | - | - | - | - | 0.684 | 0.81 | - | - | - |  | - |
| S-Adenosyl-L-homocysteine | - | - | 0.046 | 0.37 | - | - | - | - | NA | NA | - | - | - |  | - |
| N,N-Dimethylglycine | - | - | 0.003 | 0.02 | - | - | - | - | 0.307, 0.138, 0.257, 0.168, 0.517, 0.567, 0.883, 0.172, 0.125 | 0.50, 0.59, 0.47, 0.32, 1.10, 0.95, 0.34, 0.35 | - | - | - |  | - |
| Sarcosine | - | - | 0.012 | 0.12 | - | - | - | - | 0.112, 0.255 | 0.51, 0.74 | - | - | - |  | - |
| dTMP | - | - | 0.024 | 0.09 | - | - | - | - | NA | NA | - | - | - |  | - |
| Adenosine | - | - | 0.031 | 0.61 | - | - | - | - | 0.428 | 0.54 | - | - | - |  | - |
| **Glyoxylate and dicarboxylate metabolism** |  | | | | | | | | | | | | | | |
| Hydroxypyruvate | - | - | 0.008; 0.028 | 0.05; 0.07 | - | - | - | - | NA | NA | - | - | - |  | - |
| cis-Aconitate | - | - | 0.001; 0.031; 0.049 | 0.29; 0.17; 0.47 | - | - | - | - | NA | NA | - | - | - |  | - |
| L-Glutamate | - | - | 0.033 | 1.63 | - | - | - | - | 0.414 | 229.3 | - | - | - |  | - |
| 2-Hydroxy-3-oxopropanoate | - | - | 0.028 | 0.07 | - | - | - | - | NA | NA | - | - | - |  | - |
| Isocitrate | - | - | 0.002 | 0.35 | - | - | - | - | NA | NA | - | - | - |  | - |
| Pyruvate | - | - | 0.044 | 0.37 | - | - | - | - | 0.334 | 0.76 | - | - | - |  | - |
| Citrate | - | - | 0.002 | 0.35 | - | - | - | - | NA | NA | - | - | - |  | - |
| L-Glutamine | - | - | 0.011; 0.013; 0.023; 0.033; 0.044 | 1.86; 0.03; 0.007; 0.006; 0.32 | - | - | - | - | 0.485, 0.081 | 0.73, 1.60 | - | - | - |  | - |
| **Arginine biosynthesis** |  | | | | | | | | | | | | | | |
| L-Glutamate | - | - | 0.033 | 1.63 | - | - | - | - | 0.414 | 229.3 | - | - | - |  | - |
| L-Arginine | - | - | 0.042 | 0.13 | - | - | - | - | 0.295 | 1.32 | - | - | - |  | - |
| L-Glutamine | - | - | 0.011; 0.013; 0.023; 0.033; 0.044 | 1.86; 0.03; 0.007; 0.006; 0.32 | - | - | - | - | 0.485, 0.081 | 0.73, 1.60 | - | - | - |  | - |
| N-Acetyl-L-glutamate | - | - | 0.037 | 0.04 | - | - | - | - | 0.993 | 1.002 | - | - | - |  | - |
| **Nitrogen metabolism** |  | | | | | | | | | | | | | | |
| L-Glutamate | - | - | 0.033 | 1.63 | - | - | - | - | 0.414 | 229.3 | - | - | - |  | - |
| L-Glutamine | - | - | 0.044, 0.023, 0.033, 0.011, 0.013 | 0.32, 0.007, 0.006, 1.86, 0.03 | - | - | - | - | 0.485, 0.081 | 0.73, 1.60 | - | - | - |  | - |
| **Tyrosine metabolism** |  | | | | | | | | | | | | | | |
| L-Noradrenaline | - | - | 0.026; 0.002 | 0.41; 0.72 | - | - | - | - | 0.227 | 0.26 | - | - | - |  | - |
| Dopamine | - | - | 0.029 | 4.3394 | 0.04 | 0.25 | - | - | 0.253, 0.163, 0.229 | 0.06, 0.17, 0.02 | 0.179 | 0.01 | - |  |  |
| 3,4-Dihydroxymandelate | - | - | 0.038 | 0.1 | - | - | - | - | NA | NA | - | - | - |  | - |
| 3-Methoxytyramine | - | - | 0.005 | 2.78 | 0.019 | 0.63 | - | - | 0.346 | 1.31 | >0.999 | 4.75 | - |  |  |
| L-Tyrosine | - | - | 0.0008 | 0.49 | 0.028 | 0.69 | - | - | 0.85 | 1.13 | 0.643 | 1.39 | - |  |  |
| 3,4-Dihydroxyphenylacetaldehyde | - | - | 0.012 | 0.03 | - | - | - | - | 0.58, 0.226 | 0.76, 0.12 | - | - | - |  | - |
| 4-Hydroxyphenylacetaldehyde | - | - | - | - | 0.04 | 0.25 | - | - | - | - | 0.179 | 0.01 | - | - |  |
| Tyramine | - | - | 0.013; 0.037 | 2.44; 8.15 | 0.003 | 2.92 | - | - | 0.272, 0.368, 0.044 | 0.16, 0.36, 4.81 | 0.857 | 0.11 | - |  |  |
| 5,6-Dihydroxyindole | - | - | - | - | 0.028 | 0.76 | - | - | - | - | 0.286 | 40.21 | - | - |  |
| 4-Fumarylacetoacetate | - | - | 0.028 | 0.7 | - | - | - | - | NA | NA | - | - | - |  | - |
| 4-Maleylacetoacetate | - | - | 0.028 | 0.7 | - | - | - | - | NA | NA | - | - | - |  | - |
| Gentisate aldehyde | - | - | 0.047; 0.037; 0.010 | 2.02; 0.29; 0.65 | - | - | - | - | NA | NA | - | - | - |  | - |
| Pyruvate | - | - | 0.044 | 0.37 | - | - | - | - | 0.334 | 0.76 | - | - | - |  | - |
| Acetoacetate | - | - | 0.036 | 1.31 | - | - | - | - | 0.879 | 1.06 | - | - | - |  | - |
| 4-Hydroxyphenylacetate | - | - | 0.012 | 0.03 | - | - | - | - | 0.58, 0.226 | 0.76, 0.12 | - | - | - |  | - |
| **Phenylalanine metabolism** |  | | | | | | | | | | | | | | |
| Phenylacetaldehyde | - | - | 0.013 | 2.44 | 0.003 | 2.92 | - | - | 0.11 | 0.9 | 0.143 | 1.4 | - |  |  |
| -Phenylalanine | - | - | 0.026 | 0.01 | - | - | - | - | 0.169 | 2.8 | - | - | - |  | - |
| Phenylacetic acid | - | - | - | - | 0.04 | 0.25 | - | - | - | - | 0.179 | 0.01 | - | - |  |
| 2-Hydroxyphenylacetate | - | - | 0.012 | 0.035 | - | - | - | - | 0.58, 0.226 | 0.76, 0.12 | - | - | - |  | - |
| L-Tyrosine | - | - | 0.0008 | 0.49 | 0.028 | 0.69 | - | - | 0.85 | 1.13 | 0.643 | 1.39 | - |  |  |
| **Citrate cycle (TCA cycle)** |  | | | | | | | | | | | | | | |
| Isocitrate | - | - | - | - | 0.002 | 0.32 | - | - | - | - | NA | NA | - | - |  |
| Citrate | - | - | - | - | 0.002 | 0.32 | - | - | - | - | NA | NA | - | - |  |
| 2-(alpha-Hydroxyethyl)thiamine diphosphate | - | - | - | - | 0.049 | 0.05 | - | - | - | - | NA | NA | - | - |  |

^a^ P values were calculated using the Wilcoxon rank-sum test for comparisons between the S-AKI and Sepsis-only groups.

^b^ Fold change (FC) was calculated as the ratio of mean protein abundance in the S-AKI group relative to the Sepsis-only group.

^c^ Due to insufficient sample size in the Validation cohort at D8 (7 S-AKI and 2 Sepsis-only samples), Mann–Whitney U tests were performed instead of Wilcoxon rank-sum tests.

^d^ Metabolites were considered concordantly validated when the direction of fold change (up- or down-regulation in S-AKI relative to the Sepsis-only group) was consistent between the Discovery and Validation cohorts at the corresponding time point. Concordant and discordant validation results are highlighted in green and red, respectively. Metabolites assigned to multiple putative LC–MS features that exhibited both concordant and discordant fold-change directions are highlighted in yellow and excluded from the calculation of the concordance rate. Metabolites without corresponding fold-change data in the Validation cohort are highlighted in gray and treated as discordant for the calculation of the concordance rate.

^e^ – indicates that the metabolite was not identified as a pathway-enriched candidate (p < 0.05 and FC > 1.3) at the corresponding time point in the Discovery cohort and was therefore not included in the validation analysis.
